# Phase separation as a concentrating mechanism of protein/peptide hormones for the secretory granule storage

**DOI:** 10.64898/2025.12.05.692483

**Authors:** Semanti Mukherjee, Debkanya Sengupta, Prasad Sulkshane, Anindya Modak, Supriya Singh, Shouvik Manna, Ajoy Paul, Ranjit Shaw, Jyoti Devi, Shalaka Masurkar, Riya Bera, Samir K Maji

## Abstract

Protein/peptide hormones in the regulated secretory pathway are stored within secretory granules (SGs) as densely packed aggregates for extended periods of time until their stimulated release. These aggregated dense cores are reported as functional amyloids for several pituitary hormones. However, the molecular events initiating hormone aggregation and SG biogenesis remain poorly understood. Using a diverse set of protein/peptide hormones with different sizes and structures, we demonstrated that all these hormones undergo phase separation in trans-Golgi network (TGN)-relevant conditions (including low pH, the presence of glycosaminoglycans, etc.), which rapidly transitioned into solid-like structures and eventually form amyloid fibrils. Further, the cellular model expressing two protein hormones (GH-EGFP and PRL-EGFP) showed the highly dynamic, liquid-like condensate formation at the TGN volume, which was transported via microtubule to the cellular periphery/tips and formed a ready pool of mature SGs containing amyloid-rich hormone aggregates. Importantly, both in vitro and cellular studies confirmed that solid-like amyloid aggregates are reversible and can release bioactive monomeric hormones, recapitulating regulated exocytosis. Together, our findings suggest hormone phase separation at the TGN as a common molecular principle for concentrating cargo, driving rapid functional aggregation, and ensuring efficient storage within SGs.

## Introduction

All secretory proteins, including protein/peptide hormones, are synthesised in the endoplasmic reticulum (ER) and transported to the Golgi apparatus, where, from the trans-Golgi network (TGN), they are sorted into two distinct secretory pathways: constitutive and regulated secretion (1, 2). Constitutive secretion is the default pathway in all secretory cells, where proteins are continuously secreted from the cells at a rate similar to their synthesis, without any intracellular storage (3). However, specialised cells, such as neuroendocrine, exocrine, and peptidergic neurons, possess an additional pathway of regulated protein secretion, where protein/peptide hormones are sorted from the TGN to be stored within membrane-enclosed structures, called secretory granules (SGs) (4). Unlike constitutive secretion, hormones are stored inside SGs for a prolonged duration (half-life of days) at a large concentration (∼mM) (5). Upon signalling (i.e., the action of a secretagogue), these SGs are transported to the plasma membrane, fuse, and result in the burst release of hormones for their action (6). Despite extensive research, the molecular mechanism for sorting and storage of hormones inside SGs remains unclear at this point. It has been hypothesised that preferential aggregation by regulated secretory proteins segregates them from others, which might play a crucial role in sorting at the TGN and is stored as a higher-order aggregated state in SGs (7–10). In this regard, previous studies have shown that protein/peptide hormones are stored inside SGs as reversible amyloid-like aggregates (functional amyloids) (11–13), which can release the functional native proteins upon dilution at extracellular pH (11). However, it is still unknown how hormones rapidly initiate aggregation, essential for their sorting and storage during SG biogenesis and achieve a millimolar concentration regime inside SGs. Previous reports suggest that nascent SG formation initiates at the TGN as quickly as within 15-20 minutes of protein synthesis in the ER, and subsequently, the granule maturation process is completed within a few hours (14). We hypothesised that phase separation might be the key molecular mechanism for rapidly concentrating protein/peptide hormones to promote functional amyloid aggregation during SG biogenesis.

Biomolecular phase separation has emerged as a fundamental mechanism underlying the formation of membrane-less organelles within cells (15–18), performing diverse essential functions such as cell signalling, genomic organisation, transcription regulation, vesicular trafficking, and molecular sequestration, to name a few (19–21). Phase separation is driven by the cumulative effect of weak multivalent intermolecular interactions, such as electrostatic, hydrophobic, hydrogen bonding, cation-π, and π–π interactions arising from polypeptide sequences and amino acid patterning (22, 23). While these multivalent interactions are prevalent in natively unstructured proteins or intrinsically disordered proteins (24), making them primary candidates for phase separation, well-folded globular proteins can also interact through exposed surfaces, leading to crossing their solubility limit and forming the condensates (25, 26). Upon phase separation, the very high local protein concentration within condensates can also promote protein misfolding and liquid-to-solid state transition, which often leads to amyloid aggregation (27, 28). This aberrant aggregation from condensates has been observed for various neurodegenerative as well as other disease-related proteins, including α-Synuclein (α-Syn), Tau, and FUS, which may be linked to their pathological mechanisms (29–31). Interestingly, the functional liquid-to-solid transition from condensates, resulting in the formation of gels, amorphous, and amyloid aggregates, is also reported to perform diverse cellular functions such as the formation of Balbiani body, amyloid body and stress granule (32–34). This suggests that the cell strategically regulates the material states of condensates according to its functional requirements (35).

In this study, we hypothesised that functional amyloid aggregation associated with SG biogenesis initiates through biomolecular condensate formation and subsequent solid-like transition. In fact, using in vitro and cellular studies, we demonstrated various protein/peptide hormones form condensates in the presence of TGN and SG-relevant microenvironments, such as at low pH and the presence of glycosaminoglycans (GAGs). Interestingly, immediately upon condensate formation, they rapidly transitioned into a solid-like state, leading to subsequent amyloid formation in the granule-relevant conditions. Consistent with the in vitro study, the cellular study with two full-length protein hormones, GH and PRL, showed two major populations of condensates: highly dynamic condensates at the vicinity of TGN, and relatively less dynamic condensates at the periphery/tip of the cells, as suggested by super-resolution microscopy and single particle tracking. The cellular characterisation, along with other biophysical studies, demonstrated that the nascent condensates at TGN gradually matured into SGs, containing amyloid-like aggregates, which were transported to the cellular periphery/tip. Interestingly, conditions that inhibit aggregation, such as low temperature and hydrophobic stretch deletion, have been shown to inhibit phase separation/solid transition and SG biogenesis in cells. Importantly, both in vitro and in-cell hormone aggregates from condensate are capable of releasing functional native hormones under exocytosis conditions. The present study has broad implications not only in understanding the hormone storage and secretion but also in extending our understanding of phase separation-mediated amyloidogenesis in the physiological contexts.

## Results

### Phase separation of protein/peptide hormones under regulated secretory pathway-relevant in vitro conditions

To probe our hypothesis that phase separation can serve as a preferred mechanism for concentrating protein/peptide hormones during SG-biogenesis, we first selected a set of ten hormones with diverse origins, functions, and sequence lengths (**Fig. 1A, Table S1**). The secondary structure of these hormones ranges from random coil to well-folded helix, as shown by CD (**Fig. S1A)** and FTIR studies (**Fig. S1B)**. Further, in silico analysis suggests varying propensities for phase separation when analysed for the intrinsically disordered region (36) and droplet-forming potential (**Fig. S2)** (37). To examine the in vitro phase separation, all peptides/protein hormones (labelled with rhodamine, labelled: unlabelled 1:100 ratio) were studied with a 2mg/mL concentration (previously used for functional amyloid formation (11)) in 20 mM phosphate buffer at two different pH levels (cytoplasmic pH 7.4 and TGN-specific pH 6) **(Fig. 1B, C).** While none of them underwent phase separation at pH 7.4, a subset of hormones showed phase separation at pH 6. Interestingly, in the presence of other SG-associated factors, including molecular crowding and/or GAGs, all the hormones showed condensate formation in vitro **(Fig. 1C, Table S2)**.

**Figure 1:**
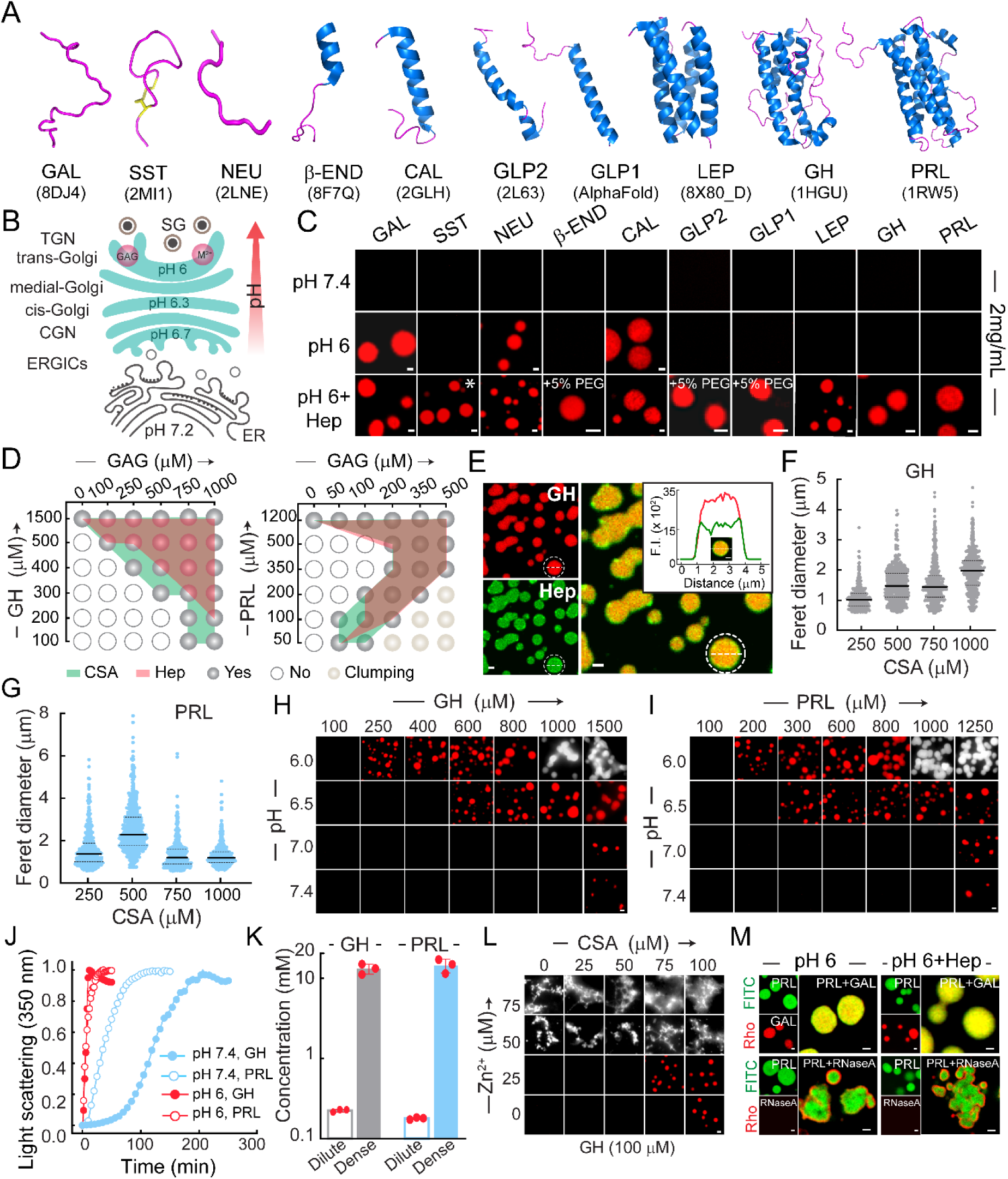
Biomolecular phase separation of a diverse set of protein/peptide hormones: **(A)** Three-dimensional structure of protein/peptide hormones as per their PDB ID, constructed in PyMOL. The GLP1 structure is predicted by AlphaFold, as the PDB structure is not available. **(B)** Schematic illustrating microenvironment/factors important for the regulated secretory pathway: low pH, metal ions and GAGs. **(C)** Condensate formation by protein/peptide hormones under different experimental conditions. *50 µM heparin for SST. **(D)** Schematic representation of GH (left) and PRL (right) phase regimes in the presence of Hep (pink) and CSA (green), showing the concentration-dependent influence of GAGs in GH and PRL phase separation. **(E)** Representative confocal microscopy images, showing the distribution of Hep within GH condensates. The inset showing the intensity profile plot of Hep(green) with GH (red) across a single GH condensate. (**F, G)** Feret diameter of GH (F) and PRL (G) condensates with varying CSA concentrations (N=550 condensates). **(H, I)** Phase regimes of GH (H) and PRL (I) with decreasing pH, indicating that low pH promotes hormone phase separation. **(J)** Static light scattering (at 350 nm) showing immediate phase separation of GH and PRL at low pH in comparison to pH 7.4 (with CSA). **(K)** Bar graph representing the dense phase and dilute phase concentration of GH and PRL. The plot represents the mean with SD, for n=3 independent experiments. **(L)** Phase regime demonstrating the effect of Zn^2+^ and CSA with 100 µM GH, where the proteins otherwise do not phase separate. **(M)** Confocal image showing multi-component phase separation of PRL and GAL (PRL+GAL) in pH 6 and pH 6+Hep, along with the individual phase separation images in the respective conditions (top row). Confocal images showing no phase separation of RNase A, along with no recruitment in PRL condensates in these two conditions (bottom row). All scale bars: 1 µm.

Further, we thoroughly investigated the phase separation property of two protein hormones (GH and PRL) **(Fig. S3)** (38). The phase regime at pH 7.4 demonstrated the essential role of GAGs in driving GH/PRL phase separation **(Fig. 1D)**. However, various GAGs (Chondroitin Sulphate A (CSA), Chondroitin Sulphate B (CSB), and Heparin (Hep)) promote a similar extent of hormone condensate formation, suggesting GAGs might act as a template for concentrating protein hormones to facilitate their phase separation **(Fig. S4)** (39). Indeed, the partitioning of fluorescently labelled Heparin into GH/PRL condensates confirmed their direct participation in condensate formation by hormones **(Fig. 1E, Fig. S5A, B)**. Further, the varied feret diameters of GH and PRL condensates in the presence of GAGs suggest that distinct GAG-protein interactions give rise to hormone-specific condensate properties upon phase separation (**Fig. 1F, G)**.

Interestingly, for these two protein hormones, when the pH of the reaction decreased gradually, we observed that lower pH levels promote their phase separation, even in the absence of GAGs **(Fig. 1H, I)**. This is further supported by condensate formation kinetics using static light scattering (350 nm), which showed a faster phase separation at pH 6 compared to pH 7.4 (even in the presence of GAGs) **(Fig. 1J)**. Further, while examining the dense/dilute phase concentration (see Methods), we observed that the dense phase concentration reached more than 50-fold higher as compared to the dilute phase, with ∼13 mM and ∼14 mM protein concentrations for GH and PRL, respectively **(Fig. 1K)**. This is indeed the cellular concentration range of densely packed hormones inside SGs (14, 40). Important to note that, along with low pH, other environmental factors, such as GAGs, metal ions, are important for SG biogenesis (**Fig. 1B, Schematic**) (8). Consistent with this, the addition of GAGs and metal ion Zn²⁺ (an abundant metal ion in SGs) at pH 6 further promoted the GH and PRL phase separation to a lower concentration regime **(Fig. 1L, S5C)**. Although GH showed strong Zn^2+^- induced clumping compared to PRL, suggesting the impact of each factor might vary for specific hormones.

To explore whether phase separation can function as a sorting mechanism during SG biogenesis, we examined the interaction of PRL condensates with a partner hormone (GAL), which is known to be co-stored with PRL (41), and with a constitutively secreted protein, Ribonuclease A(RNase A) (42). We observed selective recruitment of GAL into PRL condensates, whereas RNase A neither underwent phase separation under SG-relevant conditions (pH 6 or pH 6 with Heparin) nor was recruited into PRL condensates (**Fig. 1M**). Instead, RNase A remained on the surface of PRL hormone condensates. These findings suggest that phase-separated hormone condensates can exclude constitutive secretory proteins while selectively interacting with partner hormones, thereby enabling multicomponent phase separation relevant for their co-storage (41).

### Rapid liquid-to-solid transition of protein/peptide hormone condensates

Since the functional aggregation of hormones has been reported to be associated with SG biogenesis (7–9), we hypothesised that condensate formation of protein/peptide hormones might be associated with a liquid-to-solid transition leading to amyloid aggregation, as observed for many amyloidogenic proteins/peptides (27, 28). To probe this, we examined the macroscopic material properties of the hormone condensates under their respective phase separation conditions over time (**Table S2**). Confocal microscopy data suggested that only a subset of hormones, GAL, NEU, SST, CAL, GH, and PRL, exhibited liquid-like fusion (**Fig. 2A, Supplementary Movie 1, 2)** and surface wetting (**Fig. 2B)** immediately after formation under certain phase-separating conditions (**Table S2**). The majority of hormone condensates, however, showed neither fusion nor surface wetting even immediately after phase separation, indicating a solid-like microenvironment **(Table S2)**. The translational dynamics of molecules inside the condensates were further probed using fluorescence recovery after photobleaching (FRAP) (43). Our data revealed that the condensates of GAL, CAL, and NEU formed in the absence of GAGs at pH 6 exhibited moderate to high FRAP recovery immediately after phase separation; however, the recovery dropped rapidly in one hour (mobile fractions ∼0.2) (**Fig. 2C, D, S6A)**. Moreover, the condensates from most hormones (except β-END and NEU), formed in the presence of Heparin, displayed low initial recovery (mobile fraction ∼0.2–0.4), which further declined to below 0.2 within one hour (**Fig. 2E, F, S6B)**. These observations suggest that phase separation accompanied by rapid liquid-to-solid transition might be a shared property of regulated secretory hormones, underlining their concentrated storage inside SGs.

**Figure 2:**
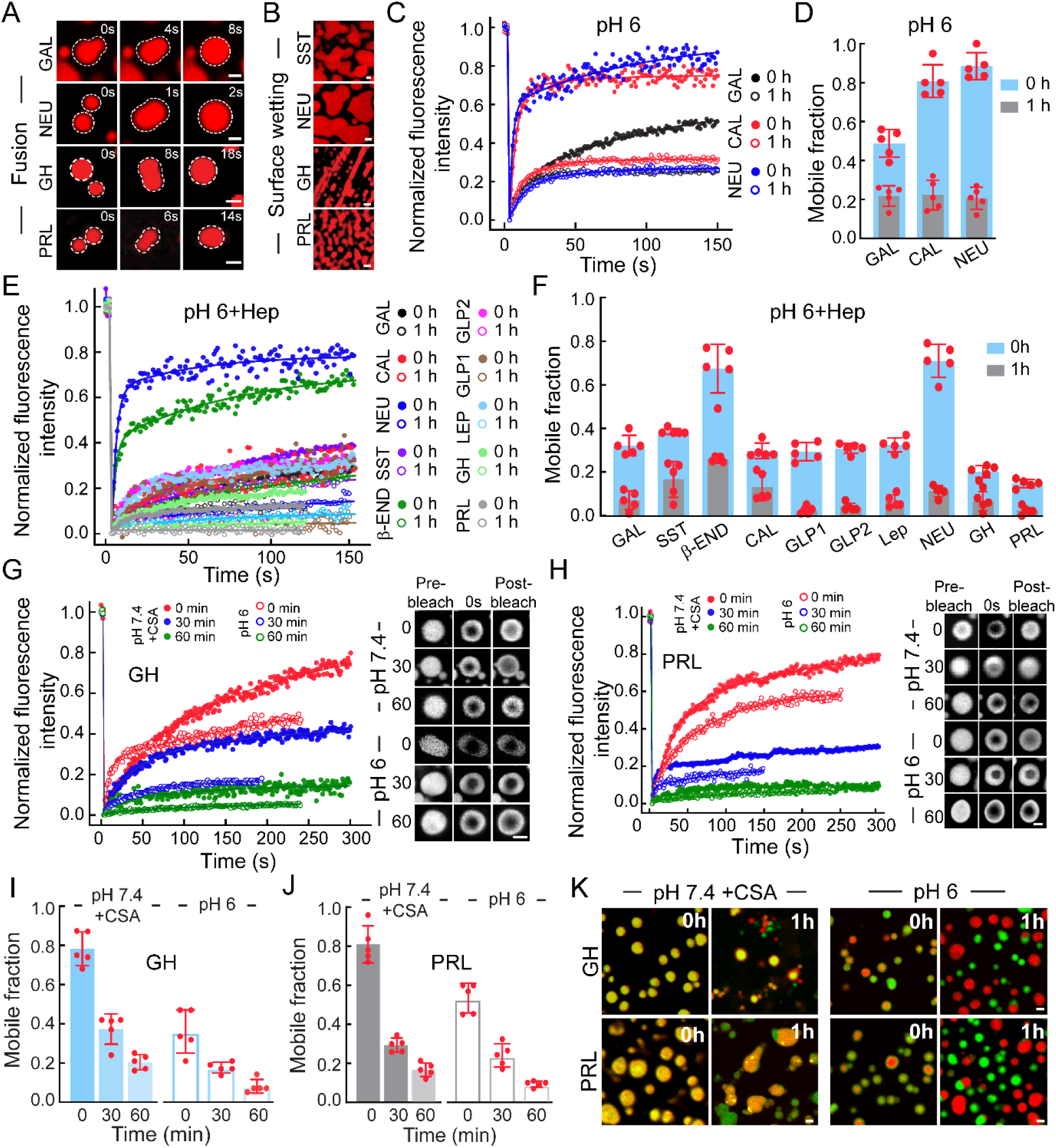
Liquid-to-solid transition of protein/peptide hormone condensates: **(A)** Fusion of two condensates immediately after their formation by GAL (pH 6), NEU (pH 6), GH (pH 7.4 + CSA), and PRL (pH 7.4 + CSA), demonstrating liquid-like properties. (**B)** Surface wetting property for SST (pH 6+Hep), NEU (pH 6), GH (pH 7.4+CSA), and PRL (pH 7.4+CSA), showing the liquid-like nature immediately after formation. (**C, D)** Normalised FRAP profiles (C) and corresponding mobile fractions calculated from five FRAP profiles for GAL, CAL, and NEU peptide condensates immediately after their formation (0 h) and after 1 h of incubation at pH 6. (**E, F)** Normalised FRAP profiles (E) and mobile fractions (F) for all hormones examined at pH 6 in the presence of Heparin, measured at 0 h and after 1 h of incubation. Bar graphs represent mean ± SD from n = 5 FRAP profiles. (**G, H**) Time-dependent (0 min, 30 min and 60 min) FRAP profiles (left) along with corresponding grayscale representative images (right) showing reduced FRAP recovery, indicating a liquid-to-solid transition of GH (G) and PRL (H) at pH 7.4+CSA (filled circle) and at pH 6 (open circle). **(I, J)** The bar graph representing the time-dependent (0 min, 30 min, 60 min) mobile fraction calculated from five different FRAP profiles for GH (I) and PRL (J), at pH 7.4+CSA (filled bar) and pH 6 (open bar), showing time-dependent reduction in the FRAP recovery. (**K)** Condensate mixing assay for GH (top) and PRL (bottom) formed under the same experimental conditions but labelled with either FITC (green) or NHS– rhodamine (red). Mixing of condensates immediately after their formation at pH 7.4 in the presence of CSA results in yellow condensates, indicating complete molecular exchange. The mixing after 1 h of incubation, as well as at pH 6 (immediately after formation), revealed distinct red and green condensates, indicating restricted exchange and reduced liquidity. All scale bars: 1 µm.

The liquid-to-solid transition of GH and PRL condensates is also promoted by SG biogenesis-associated conditions, as evidenced by their low FRAP recovery when formed at acidic pH (6.0) (**Fig. 2G-J**), in the presence of GAGs (e.g., CSA) or in the presence of metal ions (Zn²⁺) (**Fig. S6C-E**). The direct evidence of solidification is further probed by the time-dependent condensate mixing experiments, in which FITC (green) labelled preformed GH/PRL condensates were mixed with the respective NHS-Rhodamine (red)-labelled condensates (prepared separately). Mixing of two different coloured condensates, immediately after formation (0 h) at pH 7.4, resulted in a homogeneous yellow population (**Fig. 2K**). However, condensates incubated for one hour before mixing yielded distinct red and green condensates along with yellow populations. At pH 6.0, condensates mixed immediately after formation showed heterogeneous red/green distributions with only a few yellow condensates. Whereas after one hour of incubation, no yellow population was observed, suggesting rapid solidification under acidic conditions (**Fig. 2K**). Together, these observations demonstrate that rapid liquid-to-solid transition is a prominent shared feature of hormone condensates in SG biogenesis-related microenvironment.

### Amyloid aggregation from solid-like hormone condensates

We next investigated whether the solid-like hormone condensates promote amyloid aggregation, given the reported amyloid nature of several hormone aggregates associated with SG storage (11–13). The Thioflavin-S (ThS) co-partitioning, along with prominent ThS fluorescence enhancement after 12 h of condensate formation by most of the hormones under confocal microscopy, suggested that condensate solidification is associated with amyloid formation **(Fig. 3A)**. Moreover, the ThS fluorescence intensity quantification showed an increase in fluorescence at 12h; however, with hormone-specific differences **(Fig. 3B)**. This difference in ThS enhancement could be due to the extent of amyloid maturation and structural diversity of amyloids in solid condensates. The transmission electron microscopy (TEM) images further support the assumptions, revealing that although at 0h, all condensates were mostly round in morphology, after 12h of incubation, the majority of condensates showed fibril-like structures **(Fig. 3C, S7)** and amyloid-like secondary structures **(Fig. S8)**. Interestingly, some hormones (GAL, CAL, SST, β-END) showed a distinct star-shaped fibrillar outburst structure (both in ThS and EM images), well-known for the liquid-to-solid transition linked with amyloid aggregation **(Fig. 3D)** (29, 31). Further, time-dependent tracking of ThS fluorescence enhancement for selected peptide hormone condensates (GAL, CAL, and β-END) revealed that amyloid formation started immediately after condensate assembly, accompanied by the appearance of fibrillar structures, demonstrating extremely rapid aggregation kinetics (**Fig. 3E, F**).

**Figure 3:**
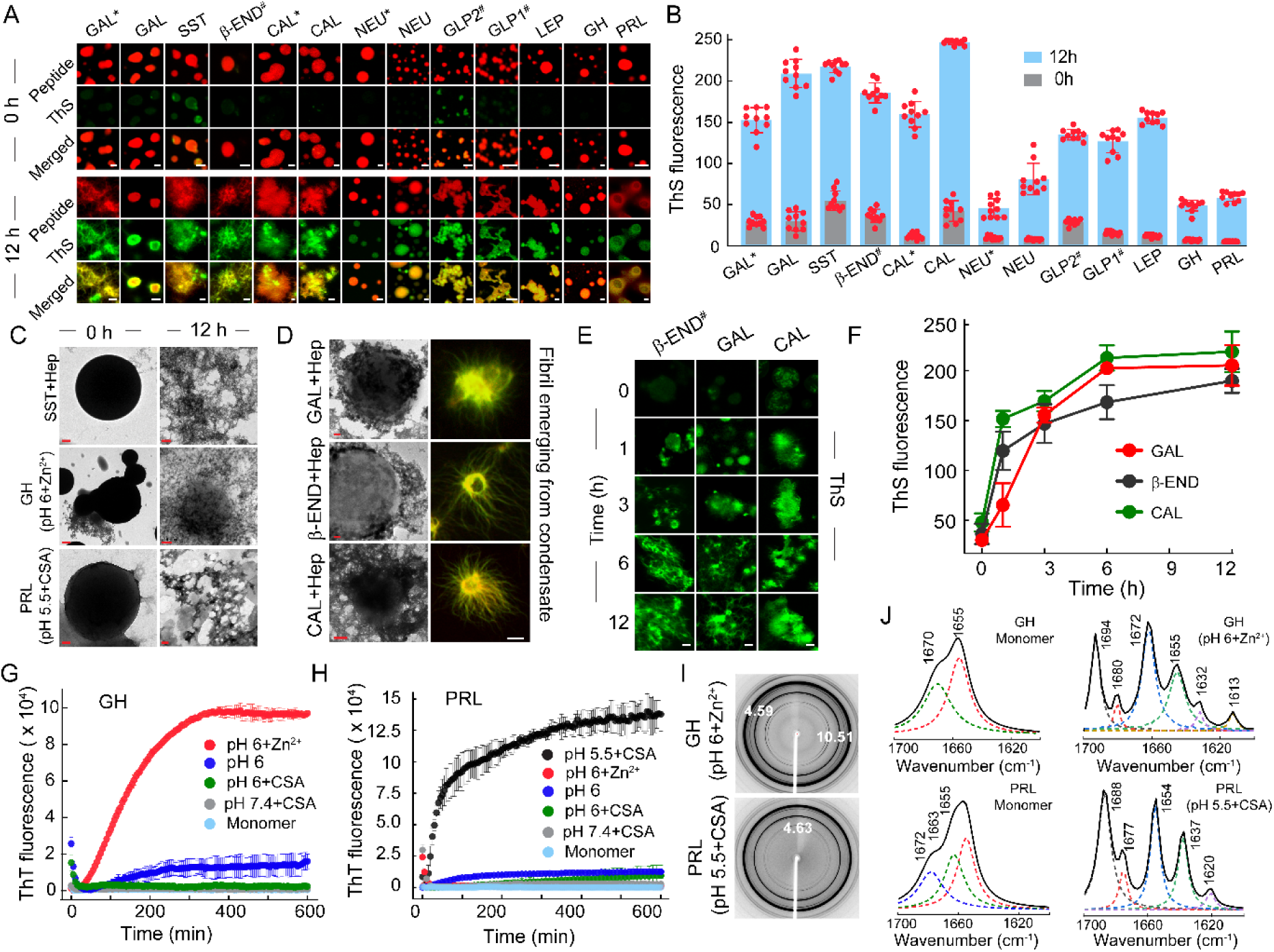
Amyloidogenic nature of the hormone condensates: **(A)** Representative confocal images showing formation of NHS-Rhodamine-labelled peptide/protein hormone condensates (red) and their binding to ThS (green). The merged profile showing ThS binding of ten different hormones at 0 h and 12h was formed under various experimental conditions at pH 6. (All conditions represent condensate formation with 500 µM Hep except *pH 6, #addition of 5% PEG-8000). (**B)** Bar plot showing the extent of ThS fluorescence increase by all hormone condensates at 0 h and 12 h. (**C)** TEM images showing spherical condensate immediately after phase separation and fibrillar morphology after 12 h of incubation, indicating that phase-separated condensates produce amyloid aggregates over time. (**D)**TEM images (left) and confocal merged images of ThS staining (right) showing fibrillar outbursts from solid-like condensates. (**E-F)** Time-dependent ThS fluorescence β-END, GAL, and CAL image and corresponding intensity calculation (F) demonstrating amyloid aggregation associated with condensate formation and solidification. (**G-H**) ThT aggregation kinetics for GH (G) and PRL (H) under various phase-separating conditions, showing higher extent of amyloid aggregation triggered by the specific granule-relevant environmental factors (Zn^2+^ + pH 6 for GH; CSA+ pH 5.5 for PRL). **(I)** X-ray diffraction of GH condensate after 24 h showing amyloid-specific diffraction patterns. **(J)** FTIR spectra of monomeric GH (top) and PRL (bottom) in comparison to their corresponding amyloid-rich solid condensates, revealing secondary structural transitions consistent with amyloid formation. Scale bars: 100 nm for TEM images; 1 μm for confocal images.

Due to low ThS signals enhancement for GH and PRL in the presence of Heparin, we examined whether any other specific conditions/factors might promote a higher amount of amyloid formation. To achieve this, we performed traditional amyloid aggregation kinetics using Thioflavin T (ThT) under various phase-separation conditions (**Fig. 3G, H**). The data revealed that GH and PRL exhibited fast and higher amyloid aggregation at pH 6 in the presence of Zn²⁺ (specific to GH SGs) and at pH 5.5 in the presence of CSA (specific to PRL SG) conditions (11, 44), although both the hormones formed condensates and subsequent solidification in several other conditions **(Fig. S9A-D)**. The amyloid formation of GH and PRL, in these SG-specific conditions, was further confirmed by X-ray diffraction pattern (reflections at 4.59 Å (equatorial) and 10.5 Å (meridional)) **(Fig. 3I)**, ThS binding **(Fig. S10)**, FTIR spectroscopy (secondary structure) (**Fig. 3J)** and TEM (morphology) **(Fig. 3C)**. Together, these results establish that solid-like hormone condensates can transition into amyloid fibrils under granule-specific microenvironments, marking a potential key step toward dense-core mature SG formation.

### Phase separation of GH and PRL forming SGs in AtT-20 cells

We further examined in-cell phase separation and functional SG formation for GH and PRL in neuroendocrine AtT-20 cells, which are known to possess the regulated secretory pathway (45, 46) **(Fig. S11)**. The immunofluorescence study showed the presence of SGs containing adrenocorticotropic hormone (ACTH) and β-endorphin (β-END), which co-localised with each other as well as with the granule matrix protein chromogranin A (CgA), typical characteristics of AtT-20 cells **(Fig. S11B-D)** (47). We transiently transfected AtT-20 cells with GH-EGFP and PRL-EGFP constructs to monitor the hormone expression via EGFP fluorescence. Previous studies have shown that the exogenous expression of a constitutive secretory protein can be rerouted in the regulatory secretory pathway into SG when fused with an SG-forming hormone component (48). Indeed, confocal imaging on post-transfected AtT-20 cells suggested a distinct appearance of hormone condensates for both GH and PRL at 16 h onwards, which are highly dynamic in nature **(Fig. 4A, S12A)**. Interestingly, the EGFP-alone construct displayed diffuse pan-cellular expression, suggesting condensate formations are due to GH and PRL **(Fig. S11E)**. Moreover, these condensate formations are specific to neuroendocrine cells, as transfection of GH-EGFP and PRL-EGFP in fibroblast cells (NIH3T3) failed to show any visible condensate formation **(Fig. S11F)**.

**Figure 4:**
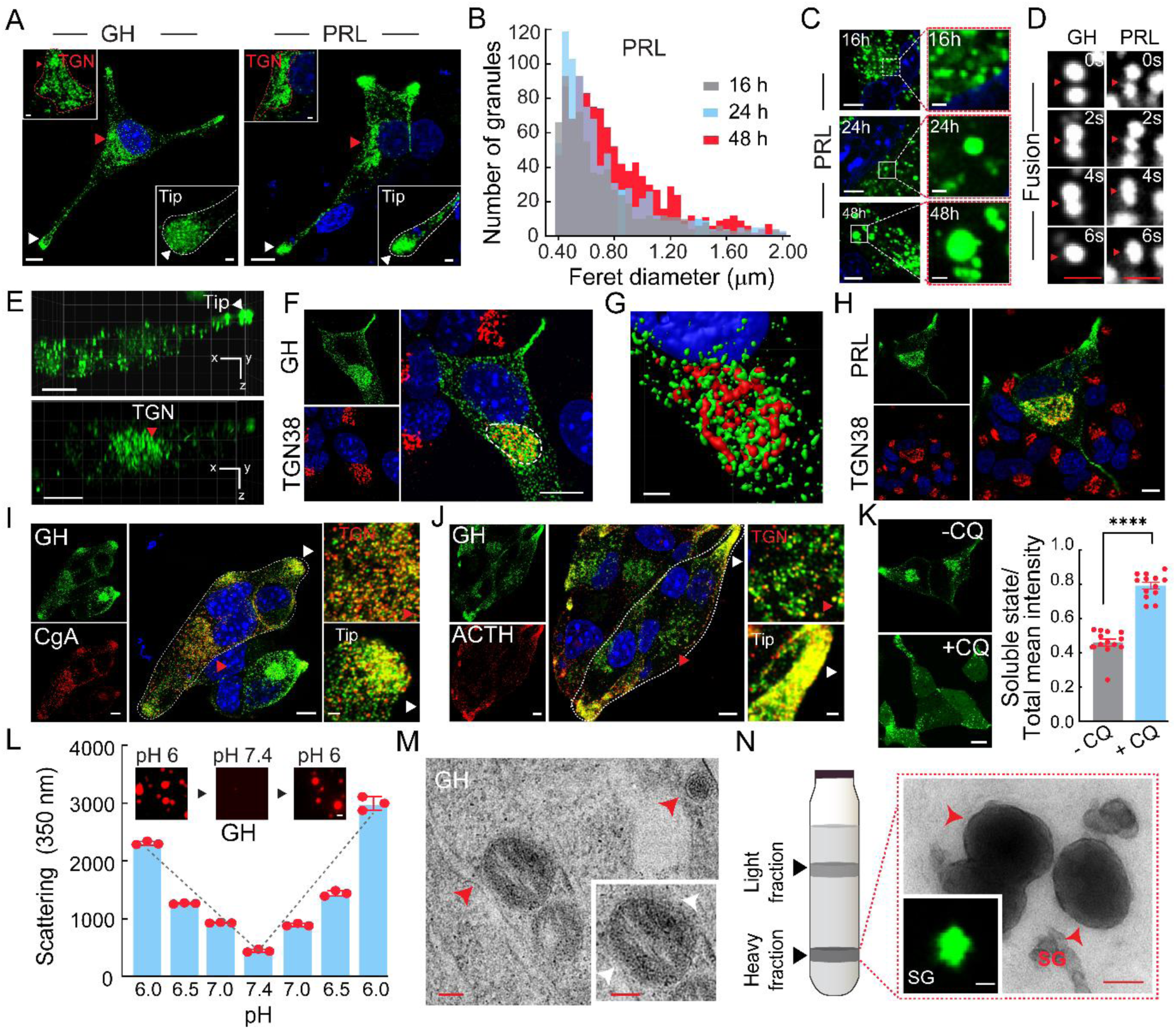
Condensate formation of GH and PRL in AtT-20 cells for their SG biogenesis and storage: **(A)** Exogenous expression of GH-EGFP and PRL-EGFP in AtT-20 cells showing two distinct populations of condensates (juxtanuclear location and the cellular tip), shown with arrows. Scale bar: 10 µm. The inset shows the magnified image of each population. Scale bar: 2 μm. **(B, C)** The feret diameter distribution (B) and corresponding confocal images (C) of PRL condensates (*n* = 10 cells) at 16 h, 24 h, and 48 h post-transfection. showing an increase in condensate size with time. **(D)** The time-lapse confocal images showing the liquid-like fusion behaviour of GH (left) and PRL (right) condensates at the juxtanuclear population. Scale bar: 1 μm. (**E)** The lattice light sheet microscopy showing GH–EGFP condensates at the tip (top) and juxtanuclear (bottom) population in the 3D volume. (**F-H)** Immunostaining on transfected AtT-20 cells showing colocalization of GH (F, G) and PRL (H) condensates with TGN antibody (TGN38). The rendering image showing volume overlap between juxtanuclear condensates (green) and TGN (red) (G). Scale bar:10 µm. **(I, J)** Immunostaining on GH- EGFP-transfected AtT-20 cells with CgA antibody (I) and ACTH antibody (J) showing colocalization of endogenous CgA, and ACTH with GH-EGFP condensates (left). Scale bar: 10 μm. Zoomed images further highlighting the colocalization at the TGN and Tip population (right). Scale bar: 2 μm. **(K)** Live cell imaging of AtT-20 cells (left) and corresponding EGFP intensity calculation (right) showing pan cellular distribution of GH-EGFP with a reduction in juxtanuclear condensates upon chloroquine treatment for 2 h. Scale bar: 10 µm. *n* = 13 cells. Statistical analysis: two-tailed unpaired *t*-test; ****p < 0.0001; 95% confidence interval. **(L)** The pH-responsive reversibility of GH condensates after solidification (at pH 6) showing dissolution of condensates upon the change in pH to 7.4. **(M)** Immuno- EM microscopy demonstrating GH-EGFP SGs (red arrow in AtT-20 cells immunolabeled with 10-nm gold particles conjugated with secondary antibody against EGFP. Scale bar: 200 nm. **(N)** (Left) Schematic showing SG isolated by density-gradient centrifugation, yielding a light granule fraction (top) and a heavy granule fraction containing dense-core SGs (bottom). (Right) TEM images of the immunoprecipitated heavy fraction containing SGs, which gives an EGFP signal under confocal microscopy (inset). Scale bars: 1 μm (confocal) and 100 nm (TEM).

Interestingly, the expression of GH/PRL-EGFP in AtT20 cells exhibited two spatially distinct condensate population: one concentrated at the juxtanuclear region and the other located at the cell periphery/tip **(Fig. 4A)**. Although these two-population remained consistent without major shift in the overall size distribution across all time points (16 h, 24 h, and 48 h post transfection) of confocal study **(Fig. 4B, C, S12B)**, however, with the peripheral population becoming more prominent after 24 h (**Fig. S12A**). Importantly, the condensates located at the juxtanuclear region are highly dynamic in nature and undergo fusion events, as shown for 24h, suggesting their liquid-like nature (**Fig. 4D, supplementary movie 3**), which is further confirmed by lattice light sheet microscopy in 3D volume (**Fig. 4E, supplementary movie 4**). In contrast, a less dynamic condensate population at the periphery/tip of the cells was observed in lattice light sheet microscopy (**Fig. 4E, supplementary movie 5**). This suggests liquid condensate formation of GH/PRL at the juxtanuclear region, undergoing fusion and partial solidification, might be located in the TGN towards forming nascent SGs, while the less dynamic population at the cell periphery might represent the storage pool of mature SGs (49–52). In fact, when we performed immunostaining with a TGN-specific antibody, TGN38 (53), confocal imaging data suggested that the juxtanuclear population (at 30h) colocalised with TGN volume, which is also confirmed by 3D volume mask rendering **(Fig. 4F-H)**. Furthermore, as expected, these condensates were neither associated with mitochondria nor lysosomes, as GH condensates in AtT20 cells did not colocalise with MitoTracker **(Fig. S13A)** and LysoTracker dyes **(Fig. S13B)**.

The SG-specific feature in GH-EGFP condensate was further supported by colocalization of CgA (a key granule matrix protein (54)) within GH condensates, both at the TGN and the cellular tip population, suggesting co-storage of CgA with GH inside SGs (**Fig. 4I**). This is consistent with the colocalization of GH and CgA in rat pituitary tissue (**Fig. S14A**). Similar colocalization with GH condensates was also observed for the endogenous secretory hormones ACTH (**Fig. 4J)** and β-END (**Fig. S14C**) in AtT20 cells, consistent with GH and ACTH colocalization in pituitary tissue (**Fig. S14B**).

We further studied whether an increase in intracellular pH impacted condensate formation and SG biogenesis of GH. In this context, membrane-permeant base chloroquine has been reported to inhibit SG biogenesis (55) and divert hormones to the constitutive pathway of secretion (56). Indeed, treating the AtT-20 cells overexpressing GH-EGFP (at 30 h) with chloroquine (200 µM) for 2h resulted in the mostly pan-cellular diffuse signal with few nascent GH condensates, as evident from confocal imaging and quantification of the condensate-to-cytoplasmic signal intensity **(Fig. 4K)**. This pH-mediated inhibition of condensate formation was further corroborated by our in vitro observations, wherein solid-like GH condensates formed at pH 6 rapidly dissolved upon increasing the pH to 7.4 and reappeared at pH 6 (**Fig. 4L**).

For direct demonstration that GH-EGFP condensates are indeed leading to SG biogenesis, we examined the SGs in the cross-sectioned AtT-20 cells through electron microscopy (**Fig. S14D**) and immuno-electron microscopy (using EGFP-specific primary antibody and secondary antibody conjugated with 10 nm gold) (**Fig. 4M**). The data showed dense-core, electron-opaque SGs that were immunolabeled with the EGFP antibody, indicating the presence of GH-EGFP-specific SGs in cells. Further isolation of SGs from cells using the previously established protocol (11) followed by immunoprecipitation using EGFP antibody revealed the membrane-bound electron-opaque dense core SG, which are rich in GH-EGFP granules, as shown by the confocal microscopy imaging (**Fig. 4N)**, EGFP and Tryptophan fluorescence in spectroscopic measurement (**S15A, B**). Together, these observations suggest that GH/PRL condensates exhibit SG-specific characteristics during their maturation, ultimately forming dense-core mature SGs.

### Condensate maturation and aggregation of hormones towards SG biogenesis

In regulated secretory cells, mature SGs containing dense-core hormone aggregates are stored at the cellular tip as a readily releasable pool for regulated exocytosis (57). We hypothesised that phase-separated GH/PRL condensates at the TGN undergo a liquid-to-solid transition while budding off from TGN as premature SGs, followed by their transport toward the cellular tip via microtubule-based trafficking (58). During this process, further cellular processing (including lowering pHs, enzymatic cleavage of pre-hormone, etc.) leads to mature, amyloid- like dense-core secretory granules (DCSGs) at the cell periphery (59) **(Fig. S20)**.

Indeed, treating the AtT-20 cells with nocodazole (depolymerising microtubules) resulted in a random distribution of GH-EGFP condensates in cells, in contrast to untreated cells (**Fig. 5A**). Further, time-dependent condensate solidification and hormone aggregation were probed by Lubrol solubility to elucidate the progression of granule maturation (60). Interestingly, Western blot analysis of different post-transfection time points (16 h, 24 h, and 48 h) revealed a progressive decrease in the soluble fraction (S), accompanied by an increase in the insoluble fraction (P) (**Fig. 5B**). The trend is consistent with the emergence of a distinct tip-localised mature SG population over time, suggesting condensate population at tip becomes enriched in aggregated hormones (**Fig. S12A**). In fact, conditions that reduce the aggregation propensity of hormone, such as decrease in temperature (20°C), inhibited maturation of SGs as GH condensates were primarily accumulated at the TGN with no detectable tip population (**Fig. 5C**). Moreover, this also resulted in a significant decrease in the Lubrol insoluble fraction of GH at 20 °C in comparison to 37 °C, confirming that the mature SG population at the tip indeed comprises hormone aggregates (**Fig. 5D**).

**Figure 5:**
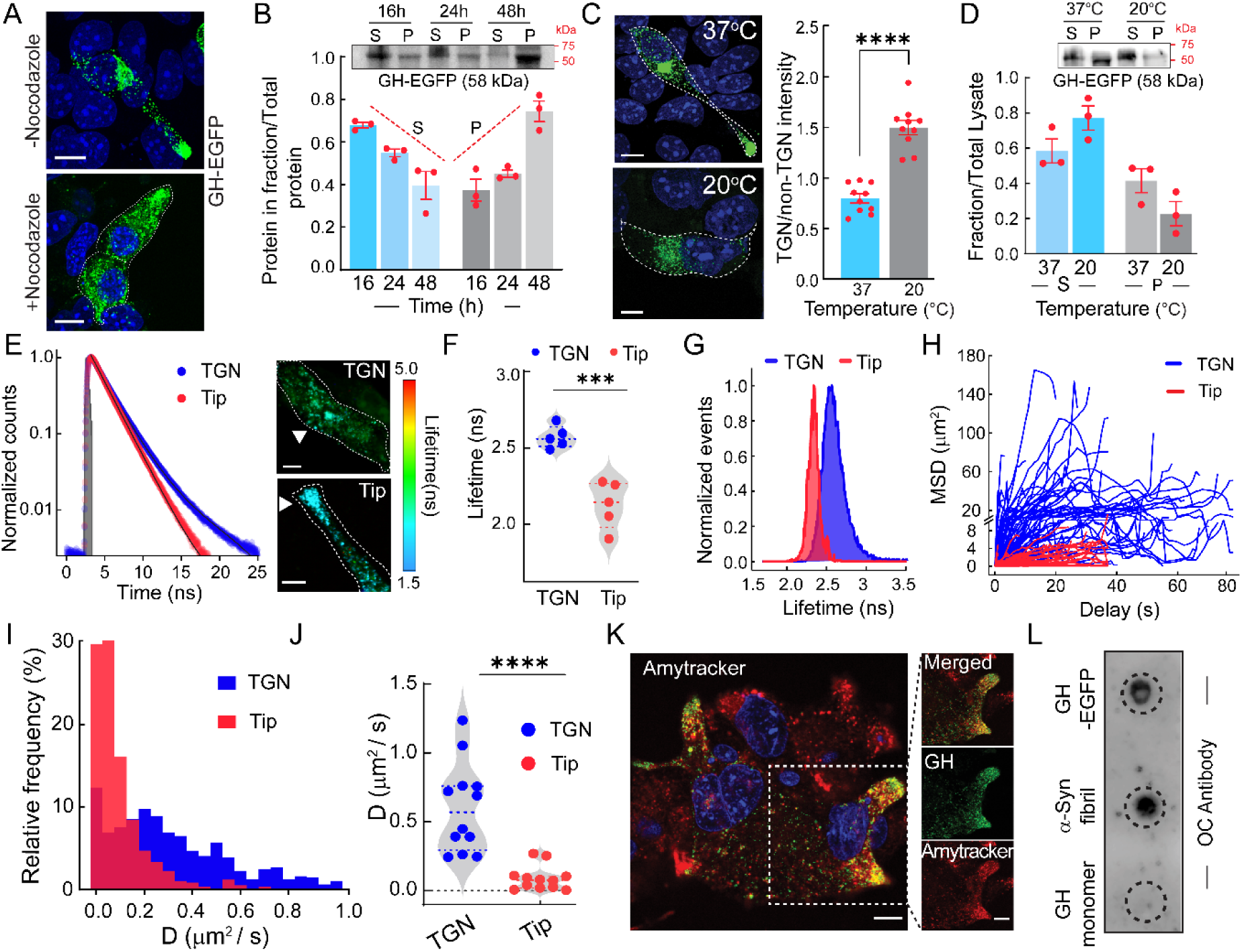
Hormone aggregation leading to maturation of SGs: **(A)** Treatment with nocodazole showing disruption of GH condensates/granules localization in AtT-20 cells. Nocodazole untreated cells were used as a control. Scale bar: 10 µm. **(B)** Western blot showing the time-dependent (16 h, 24 h, and 48 h post-transfection) increase in Lubrol-insoluble (P) fractions of the GH-EGFP population. S and P representing the Lubrol soluble and insoluble fraction, respectively (top). Representing the quantification of S/total and P/total fractions from three independent blots, demonstrating a progressive increase in the insoluble fraction and a corresponding decrease in the soluble fraction over time (bottom). **(C)** Live-cell confocal imaging of GH-EGFP-expressing AtT-20 cells (left) and corresponding intensity calculation (right) showing a decrease in GH SGs at the cellular Tip when the culture temperature is reduced to 20°C. Scale bar: 10 µm. Statistical analysis using two-tailed unpaired t-test (****P ≤ 0.0001 with 95% confidence interval. **(D)** Western blot (top) and corresponding intensity calculation (bottom) showing an increase in the Lubrol-soluble fraction when cells were incubated at 20°C compared to 37°C. **(E)** Fluorescence lifetime imaging (scaled 1.5-5.0 ns, left) and corresponding lifetime decay curves (right) of GH-EGFP condensates in AtT-20 cells showing faster decay and lower lifetime at the Tip population compared to the TGN population. **(F)** Violin plot representing the average fluorescence lifetime of the TGN and Tip population (N=5 cells). Statistical analysis used a two-tailed unpaired t-test (****p<0.0001 with 95% confidence interval. (**G)** The lifetime distribution showing a broader distribution at TGN (blue) compared to Tip (red). (**H)** MSD profile of all GH-EGFP condensates/SGs showing higher displacement at TGN (blue) compared to the Tip (red). **(I, J)** Distribution of diffusion coefficient, D (I), and average D values (J) for condensates at the TGN (blue) and Tip (red). Tip condensates display lower average D values along with a narrower distribution compared to the TGN population (N = 500). Statistical analysis: two-tailed unpaired t-test (****p<0.0001 with 95% confidence interval). **(K)** Live cell confocal imaging of GH-transfected AtT- 20 cells with Amytracker dye showing the colocalization of the dye with SGs, suggesting amyloid-like hormone aggregation. Scale bars: 10 µm; 2 µm (inset). **(L)** Dot blot assay with immunoprecipitated GH-EGFP cell lysate showing OC immunoreactivity with GH-EGFP fraction, confirming their amyloid nature. α-Syn fibril and GH monomeric protein were used as positive and negative controls, respectively.

Collectively, these findings indicate distinct material properties of condensates populated at the TGN and at the cellular tip, which was further supported by fluorescence lifetime imaging microscopy (FLIM) of GH–EGFP expressing AtT-20 cells (30h) (**Fig. 5E**). The tip-localised condensates exhibited reduced fluorescence lifetimes (2.13 ± 0.15 ns) (**Fig. 5F**), a narrower lifetime distribution (**Fig. 5G**), and faster decay kinetics (**Fig. 5E**) compared to the TGN population (2.57 ± 0.07 ns), indicating a more homogeneous and aggregated state. In contrast, the TGN-localised condensates displayed broader and more heterogeneous lifetime distributions, suggesting a mixture of condensates at different stages of the liquid-to-solid transition (**Fig. 5E-G**). Consistent with this, the single-particle tracking further revealed distinct dynamic behaviours of condensates at the TGN and cellular tip. The TGN-associated nascent granules exhibited a ∼10-fold higher mean square displacement (MSD) (**Fig. 5H)** and a higher average Diffusion coefficient (D) with a heterogeneous distribution (**Fig. 5I, J)** compared to tip-localised mature condensates/SGs (**Fig. H-J, S16**).

We next examined whether the aggregates of mature SGs possess amyloid-like characteristics (11–13). Live-cell confocal imaging of GH-expressing cells stained with the amyloid-specific Amytracker 630 dye (61) showed that the GH SGs (at the tip) exhibited high colocalization with the dye, suggesting the amyloid-like feature of mature SGs (**Fig. 5K)**. This was further supported by the dot blot assay of immunoprecipitated GH–EGFP from AtT-20 cell lysates, which showed positive immuno-staining with the amyloid-specific OC antibody (62) (**Fig. 5L**). Together, these findings support the hypothesis that condensate maturation leads to hormone aggregation into amyloids, which are stored as mature SGs at the cellular tip.

### Amyloid aggregates of hormones retain functional properties

We further examined whether phase separation and condensate-mediated amyloid-like mature SGs are responsive to secretagogue-stimulated exocytosis. We observed the complete loss of GH condensates/granules from the cellular tip, under confocal microscopy, upon BaCl2 treatment (used as a secretagogue (63)) for 30 minutes in AtT-20 cells **(Fig. 6A, B).** Consistent with this, Western blot analysis and the EGFP fluorescence signal at 508 nm from the collected culture media showed a BaCl2 dose-dependent increase in the released hormone, suggesting that GH-EGFP condensates indeed formed biologically active SGs with the releasable pool of hormones **(Fig. 6C).**

**Figure 6:**
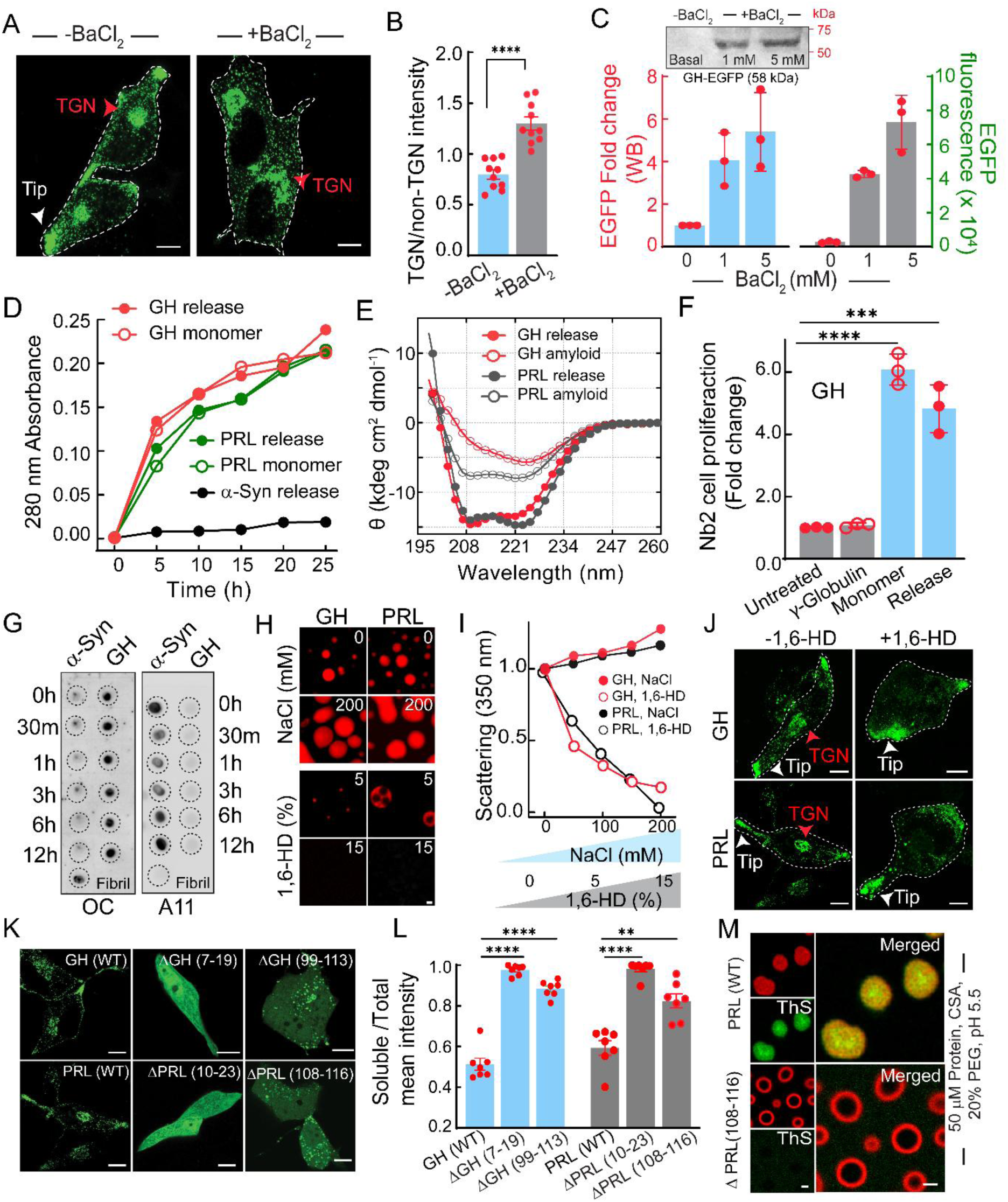
Structure-activity and reversibility of amyloid-rich hormone condensates for their SG biogenesis. (A, **B)** Live cell confocal imaging (A) and corresponding intensity calculation (B) showing absence of GH condensates/SGs at Tip population when treated with 1 mM BaCl2 (+BaCl2). Scale bar: 10 µm. Statistical analysis: two-tailed unpaired t-test (****P ≤ 0.0001 with 95% confidence interval). **(C)** Western blot analysis of cell media (top) and corresponding intensity calculation (bottom) in the absence and presence of various concentrations of BaCl2, showing dose-dependent enhancement of GH secretion into the media. The right axis representing the EGFP fluorescence at 508 nm from the released media (bottom). **(D)** Monomer release profile of GH (red) and PRL (green) amyloid-rich condensates, monitored using UV absorbance at 280 nm, showing the gradual release of hormone monomer from solid, amyloid-like condensates. α-Syn solid condensates showed no release of monomer (black) with time in an identical experimental setup. **(E)** CD spectra showing secondary structure of the released GH and PRL monomer (GH/PRL release), from the solid-like amyloid-rich condensates (GH/PRL amyloid). **(F)** Fold change in Nb2 cell proliferation in the presence and absence of released GH. The monomeric protein and untreated cells were used as a positive control and a negative control. Unrelated protein γ- Globulin also showed no cell proliferation. Statistical analysis: two-tailed unpaired t-test (****P ≤ 0.0001,***P ≤ 0.001with 95% confidence interval). **(G)** Time-dependent dot blot analysis for GH and α-Syn condensates, with amyloid-specific OC antibody (left) and oligomer-specific A11 antibody (right). **(H)** The NaCl and 1,6-Hexanediol (1,6-HD) titration followed by static light scattering experiments using preformed condensates by GH and PRL, showing dissolution of condensate with 1,6-HD treatment in contrast to NaCl. Corresponding confocal images are also shown for selected concentrations of both additives. **(J)** Live cell confocal images showing dissolution of the condensate population at TGN by GH-EGFP in AtT-20 cells after 1,6-HD treatment, Scale bar: 10 μm. **(K, L)** The confocal images (K) and corresponding EGFP intensity quantification (L) showing expression and inhibition of condensate formation by the amyloid deleted region of various constructs (GH (WT), ΔGH (Δ7-19), ΔGH(Δ99-113 a.a), ΔPRL 1(Δ10-23 a.a), ΔPRL 2(Δ108-116 a.a)). Scale bar: 10 μm. Statistical analysis: two-tailed unpaired t-test (****P ≤ 0.0001, ****P ≤ 0.0001, ****P ≤ 0.0001, **P ≤ 0.05 with 95% confidence interval). **(M)** In-vitro phase separation of ΔPRL(Δ108-116 a.a) showing hollow condensate formation, which does not bind to amyloid-specific dye ThS (bottom) in contrast to WT PRL. Scale bar: 1 µm.

The in-cell data were further supported by in vitro solid/amyloid condensate monomer release data (11), where amyloid-rich solid condensates were shown to release monomers at pH 7.4 (extracellular pH) by both the GH and PRL, as monitored through UV absorbance (**Fig. 6D)**. The released hormone not only possessed the native structures (GH/PRL), as observed in CD spectroscopy (**Fig. 6E),** but also showed functional activity as evident from Nb2 cells proliferation assay (containing GH/PRL receptor) (**Fig. 6F, S17)** (64). Important to note that, in contrast to GH/PRL solid condensates, the PD-associated α-Syn solid condensates did not release any monomer under identical conditions (**Fig. 6D)**. This suggests that reversibility and ready release of monomeric protein from solid amyloid-like condensates might be the property of functional amyloid associated with SG biogenesis (11). Apart from this, the rapid solidification of these SG hormone condensates may further reduce the probability of cytotoxic oligomer accumulation, in contrast to the solidification of disease-associated condensates. In fact, when we probed dot blot using OC (specific to fibrils) and A11 (amyloid oligomers) antibodies, we observed no A11 immunoreactivity during GH condensate formation and solidification, in contrast to α-Syn (**Fig. 6G)**. This differential property of faster solidification and release might be encoded by the protein/peptide hormone sequence and the microenvironment of condensate formation.

Further, the NaCl and 1,6-Hexanediol (1,6-HD) titration, determining the type of interaction underlying GH/PRL condensate formation (65), suggested that the weak hydrophobic interaction plays a major role in their phase separation, condensate stability and probably for further solid-like maturation (**Fig. 6H, I)**. This is further supported by the observation of rapid dissolution of nascent condensates of GH-EGFP and PRL-EGFP at the TGN of AtT-20 cells upon treatment with 1.5% 1,6-HD for 30 min, whereas mature amyloid-like aggregates at the cellular tip remained largely unaffected (**Fig. 6J**). To further examine the importance of the amyloid-forming region of the hormone for the condensate formation and solidification in vitro and in cells, we constructed various deletion mutants, ΔGH (7-19), ΔGH (99-113), ΔPRL (10- 23), ΔPRL (108-116), along with a double-deleted mutant, ΔΔPRL (10-23, 108-116) (Fig. S18A-C, Table S3, see method). Although removing the N-terminal amyloidogenic region from the signal sequence [ΔGH (7-19), ΔPRL (10-23), or double mutant ΔΔPRL(10–23,108– 116)] led to diffuse, pan-cellular distribution of protein with no condensate formation **(Fig. 6K, L, S18D, E)**, deletion of the internal aggregation-prone motifs [ ΔGH (99-113) and ΔPRL (108- 116)] alone resulted drastic reduction of GH/PRL condensates along with an increase in pan-cellular expression (**Fig. 6K**). Consistent with cellular data, the in vitro phase separation of the ΔPRL (108-116) (**Fig. S19 A-D)** revealed primarily hollow condensate formation at pH 5.5 in the presence of CSA, which neither underwent solidification nor amyloid formation as shown by FRAP recovery and ThS staining, in contrast to the WT protein (**Fig. 6M, S19E**). These cumulative observations demonstrate that amyloidogenic motifs encode the structural information necessary for condensate formation and maturation, thereby leading to the functional role in granule biogenesis.

## Discussion

SG biogenesis is a highly regulated process, essential for the storage and release of protein and peptide hormones in response to stimuli in endocrine and neuroendocrine cells (66, 67). Immature SGs are formed in the TGN by selective sorting, aggregation, and condensation of freshly produced cargo proteins (1, 4). These granules then mature through luminal acidification, membrane remodelling, and enzymatic processing, resulting in dense-core structures capable of controlled secretion (68, 69). Previous studies have suggested that aggregated protein/peptide hormones in SG are functional amyloids in nature, capable of releasing monomeric hormones in response to changes in extracellular pH or large dilution (11–13). However, the molecular factors that trigger cargo aggregation into functional amyloids and govern the initial stages of SG biogenesis remain elusive.

Phase separation is a crucial biological phenomenon that underlies the sequestration of macromolecules within membrane-less organelles in cells (15–18). Biomolecular phase separation spatiotemporally regulates diverse cellular functions, where, depending on their functional roles, condensates can exhibit a wide range of material properties, from classical liquid-like droplets (e.g., germline P granules, nucleoli, etc. (70–72)) to gel-like assemblies (etc., centrosomes, nuclear pore complexes, etc (73, 74)) and even amyloid-like functional aggregates (e.g., Balbiani bodies and amyloid bodies (33, 34)). The active cellular machinery tightly regulates the condensate’s material states and actively disassembles solid-like, non-dynamic condensates in a timely manner once their functional role is fulfilled, thereby preventing them from aberrant solidification (35). For instance, reversible gelation of yeast prion protein Sup35 condensates (75), FUS (76) and amyloid-like condensates (e.g Balbiani bodies and amyloid bodies (33, 34)) undergo regulated dissolution after completing their functions. In this study, we hypothesised that phase separation, followed by a rapid liquid-to-solid transition, might be the driving force preceding functional amyloid formation of these protein/peptide hormones during SG biogenesis (**Fig. S20**).

Upon investigating many (a total of ten) different hormones, which varied in size, length, and secondary structure, we observed that all hormones undergo phase separation under conditions conducive to SG biogenesis, including low pH, the presence of metal ions, and GAGs in an in vitro setup. However, the impact of each factor on individual hormone phase separation varied, suggesting context-dependent molecular interactions. Interestingly, all of them exhibited an instantaneous liquid-to-solid transition, leading to their aggregation; however, their amyloid growth and maturation can further depend on more specific factors, for instance, as observed for PRL (in the presence of CSA) and GH (in the presence of Zn^2+^) (11, 12). This indicates that, although hormone phase separation is initiated at the TGN, additional SG-specific triggers and/or specific partner molecules/other hormones are likely required to drive amyloid maturation from condensates to form dense-core mature SGs. Indeed, this finding is consistent with a previous study, which demonstrated that Zn²⁺ ions can trigger the formation of filamentous aggregates from chromogranin-B condensates (77). Further pre-pro- hormone/prohormone, which is reported to have less aggregation tendency (78, 79), might also form liquid-like condensates while being processed by the enzyme, which makes active hormones resulting in their high aggregation and solidification towards mature SG biogenesis (77, 80).

Upon delving deeper at the cellular level, GH/PRL-EGFP expression in AtT-20 cells resulted in intracellular condensate formation and displayed the characteristic nature of SGs, such as a pH-responsive nature, partitioning with CgA, and a membrane-enclosed granular morphology. Interestingly, these cells exhibited two distinct condensate populations, consistent with previous reports of SG localisation (49, 50). The population, localised at the TGN-associated juxtanuclear Golgi region, is highly dynamic, undergoing occasional fusion, and largely represents the Lubrol-soluble fraction. Further, their dissolution with 1,6-HD and broad distribution with higher life-time in contrast to tip population suggest the TGN population of condensates possess heterogeneous material properties, which are due to the different stages of their liquid-to-solid state transition. Further, the decrease in granule maturation at low temperature (at 20°C), as evident from the reduced tip population and the Lubrol-insoluble fraction, renders direct evidence that phase-separated nascent granules are eventually aggregated, forming the tip-localised mature granule pool. These aggregates from mature SGs exhibited an amyloid-like nature, as shown by their binding to the amyloid-specific antibody OC and the amyloid-specific dye Amytracker (**Fig. 5K**). The aggregation and amyloid maturation process may be governed by hydrophobic interactions and/or mediated by the amyloidogenic region, as both in vitro condensates (GH and PRL) and nascent intracellular condensates formed at the TGN were dissolved upon 1,6-HD treatment (**Fig. 6J**), as well as upon deletion of the amyloidogenic region from the respective protein/peptide hormones. (**Fig. 6K**).

To view the sorting of regulated versus constitutive secretory proteins through the lens of biomolecular phase separation, specific environmental factors may be necessary to enable the selective phase separation and solidification of hormones. This might exclude the constitutively secretory proteins that either lack phase separation propensity under SG conditions or have high critical concentration requirements to undergo phase separation, which may not be easily achievable due to their continuous release (2). Apart from hormone-specific environmental factors, the condensate formation and subsequent solidification process are also dependent on the amino acid sequence and structure of the protein/peptide in the physiological context, as well as their interaction with partner hormones (11, 41), which may dictate selective protein recruitment within the condensate. Although the possibility cannot be ruled out that constitutive secretory proteins may also form liquid condensates, which do not mature into stable dense-core aggregates. This may enable them to be continuously incorporated into budding vesicles that traffic directly to the plasma membrane. Indeed, we observed that, while constitutively secretory protein RNase A neither phase separated nor partitioned into PRL condensates in vitro at SG-relevant conditions, however, Galanin (known to be co-stored with PRL (41)) readily co-phase separated with PRL to form a multicomponent condensate (**Fig. 1M**).

One of the major concerns associated with amyloid formation is the potential cytotoxicity arising from oligomeric intermediates. Notably, the rapid solidification with amyloid aggregation observed for protein/peptide hormones (**Fig. 3**) suggests that SG-associated environmental cues and/or peptide sequences have been evolutionarily designed for immediate aggregation, thereby minimising the possibility of cytotoxic oligomer formation/accumulation (81), as shown for GH aggregation **(Fig. 6G)**. Furthermore, encapsulation of these aggregates within an “inert” membrane can mitigate potential toxicity while allowing for their disassembly upon exocytosis.

Moreover, the irreversible nature of amyloid architecture and its abnormal deposition, which are frequently linked to adverse pathological outcomes in several diseases, are the primary reasons for concern (82, 83). Importantly, both intracellular amyloid-like mature SGs and in vitro amyloid-rich condensates released functional hormones with the native conformation under exocytosis-mimicking conditions. This is consistent with previous reports showing that functional amyloid is responsive to pH changes (e.g., pH-dependent protonation/deprotonation of short structural motifs (13, 84, 85) and dilution (11), which might have been evolutionarily designed for the rapid aggregation/disaggregation process and to minimise amyloid-associated toxicity. However, dysregulation of such amyloid disassembly mechanisms may contribute to pathological amyloid accumulation observed in several endocrine disorders, including pituitary adenoma, type II diabetes, medullary carcinoma, and atrial amyloidosis (86–88). Thus, phase separation–mediated reversible aggregation of protein/peptide hormones into functional amyloids underscores the fine balance between physiological storage mechanisms and pathological amyloidogenesis, highlighting the tight relationship between phase separation, peptide aggregation, and function.

## Acknowledgments

The authors acknowledge the IIT Bombay central facility and SAIF for TEM, FACS, FTIR, NMR, and XRD facilities. SKM acknowledges DST SERB-SUPRA (SPR/2021/000103), JC Bose Fellowship (JCB/2023/000027), Wadhwani Research Centre for Bioengineering (WRCB), TATA Innovation (BT/HRD/35/01/03/2020), DBT (BT/PR42116/MED/32/779/2021), and Sunita Sanghi Centre for Neurodegenerative Diseases (SCAN), IIT Bombay, for financial support. S.M. acknowledges the Ministry of Education (MoE), Government of India, for the fellowship. We thank Prof. Praful Singru (National Institute of Science Education and Research) for the kind gift of tissue sections.

## Author Contribution

The study was conceptualised by S.K.M. and designed by S.K.M. and S. Mukherjee. S. Mukherjee, A.M., S.S., D.S., S. Manna, J.D., R.S., R.B. performed the in-vitro experiment. S. Mukherjee, D.S., P.S., A.P. and S. Masurkar performed in-cell experiments. S.K.M., S. Mukherjee and D.S. participated in the manuscript writing. S. Mukherjee prepared the illustration. S.K.M., S. Mukherjee, R.S., S. Manna, A.P., A.M., and D.S. edited the manuscript draft. All the authors discussed and approved the final version of the manuscript.

## Data Availability

All the data in this manuscript are available in the main and supplementary information. The tools used for plotting and analysis have been mentioned in the main and supplementary manuscripts.

