## supplementary information for "Phase separation as a concentrating mechanism of protein/peptide hormones for the secretory granule storage"

### Contents

**Page no.**


| | | |
|---|---|---|
| 1. | Materials and Methods | 3 |
| 2. | Supplementary Figures (1-20) | 25 |
| 3. | Supplementary Tables (1-3) | 45 |
| 4. | Supplementary Movie Legends (1-5) | 49 |
| 5. | References | 50 |

#### MATERIALS AND METHODS

##### Chemical and Reagent

All chemicals and reagents used in the experiments were mostly purchased from Sigma-Aldrich (St. Louis, MO, USA), Merck (Darmstadt, Germany), or HiMedia (India), unless otherwise specified. Dye used for protein/peptide labelling, NHS-Rhodamine (Catalogue no. 46406), and Fluorescein-5 isothiocyanate (FITC) (Catalogue no. F1906), were obtained from ThermoFisher Scientific (USA). PD10 column, Superdex$^{TM}$ 200 size-exclusion column, and HiTrap anion exchange column were obtained from GE Healthcare Life Sciences (U.S.A). Peptides, GAL, SST, β-END, GLP1, GLP2 and LEP were manufactured by BACHEM. Salmon calcitonin (Catalogue: 05232401), Neurotensin (Catalogue: N6383) and Ribonuclease A(Catalogue: R6513) were purchased from Sigma-Aldrich. Protease inhibitor cocktail (PIC) was procured from Roche Applied Science (Catalogue no. 05056489001). Lubrol (Catalogue no. 195299) was purchased from MP Biomedicals. Transfection reagents Lipofectamine and P3000 were purchased from Invitrogen, USA (Catalogue no L3000-008). All western blots were developed using the Bio-Rad ECL detection kit (Catalogue no. 1705060). All the buffer solutions were prepared using double-distilled and deionised water from the Milli-Q system (Millipore Corp., Bedford, MA), adjusted to pH (± 0.01) at 25°C, and filtered with a 0.22 µM filter paper.

##### Molecular cloning

Wild-type full-length GH and PRL genes (without N-terminal signal sequence) in the pT7L plasmids were kindly gifted by Prof. PS Dannies and Prof. ME Hodsdon from Yale University School of Medicine and were used for recombinant bacterial expression for in vitro experiments. Since a signal sequence is essential for protein synthesis in the mammalian system and to translocate nascent protein into the ER lumen, the signal sequence of Human GH (aa: MATGSRTSLLLAFGLLCLPWLQEGSA) was incorporated at the N terminus of the Full-length GH through molecular cloning and in the pEGFP-N1 vector, which has been used for all in-cell experiments. The cloning was performed through polymerase chain reaction (PCR) using the following forward primer: 5’-CCCAAGCTTATGGCCACCGGCAGCAGGACCAGCCTGCTGCTGGCCTTCGGCCTGCTGTGCCTGCCCTGGCTGCAGGAGGGCAGCGCCTTCCCAACCATTCCC-3’ and reverse primer 5’-CCGCTCGAGGAAGCCACAGCTGCCC-3’. A second PCR was performed with the forward primer 5’-GCTCAAGCTTGCCACCATGGCCACCGGCAG-3’ and reverse primer 5-GGTGGATCCTCGAGGAAGCCACAGCT-3’ to incorporate HindIII and BamHI restriction enzyme sites at the two ends of the GH amplicon, respectively. After the second PCR amplification, restriction enzyme digestion was performed on the GH amplicon and on the pEGFP-N1 vector, followed by the ligation reaction (T4 DNA ligase) and transformation into XL10 (Gold) ultracompetent cells. The colonies containing the plasmid were selected using the kanamycin antibiotic resistance in the Luria-Agar culture plate. The colonies were inoculated in Luria broth (LB) medium containing kanamycin antibiotic. Plasmids were isolated by using a mini-prep kit (GeneAll Exprep$^{TM}$ Plasmid SV mini), and concentration was measured using a NanoDrop UV-Vis spectrophotometer (Implen, USA). The newly cloned plasmids were verified by Sanger sequencing.

Full-length PRL (with N-terminal signal sequence) was amplified from plasmid “Prolactin-CD4d3+4-bio” (Addgene ID #73123) with the forward primer 5’-ATAAAGCTTGCCACCATGAACATCAAGGGCTCCCCA-3’ and reverse primer 5’-GGTGGATCCTCGAGACAGTTGTTGTTGTGGATG-3’ to introduce HindIII and BamHI restriction enzyme sites at the two ends of the PRL amplicon, respectively followed by Restriction enzyme digestion, ligation in pEGFP-N1 vector and plasmid isolation were performed as previously described. The final plasmid construct was verified by Sanger sequencing.

All the amyloidogenic sequences deletion variants of GH and PRL, ΔGH (7-19), ΔGH (99-113), ΔPRL1(10-23), and ΔPRL (108-116), ΔΔPRL (10-23, 108-116) were cloned in the pEGFP-N1 vector, and are synthesised by Thermo Fisher Scientific, USA and verified by Sanger sequencing.

**In silico analysis of proteins**

The secondary structure proteins/peptides were obtained from the Protein Data Bank (PDB), except for GLP1, whose structures were not deposited in the PDB and were therefore predicted using the AlphaFold Server (1). Table S1 summarises the FASTA sequence, structural information, and the PDB and UniProt IDs of all proteins/peptides. The output PDB/AlphaFold structures were represented by using PyMOL (2). The FASTA amino acid sequence of all full-length proteins and peptides was used for a range of in silico analyses. The online platform IUPred2A was used to identify the disordered region in the primary protein structure (3, 4). The FuzDrop tool was used to predict the overall phase separation propensity of protein/peptides and to identify the droplet-promoting region (DPR) (5, 6). The in silico tool, SMART (Simple Modular Architecture Research), was used to identify the low-complexity regions (LCRs) on the protein/peptide sequence (7). The amyloidogenic region of the protein/peptide primary sequence was predicted by the TANGO software (8). All the output data from in-silico analysis were plotted using OriginPro 2021 (Origin Lab, USA) software.

**Recombinant protein expression and purification of GH, PRL and ΔPRL (108-116)**

Human GH and PRL were recombinantly expressed in *Escherichia coli* BL21 (DE3) following previously reported protocols with minor modifications (9, 10). Briefly, GH, PRL, and ΔPRL (108–116), encoded in pT7L plasmids, were transformed into *E. coli* BL21 (DE3) competent cells. The transformed cells were selected on ampicillin-containing plates, and a single colony was inoculated into 100 mL LB medium (primary culture) and grown at 37°C and 200 rpm until the optical density at 600 nm ($OD_{600}$) reached ~0.6. For the secondary culture, 20 mL of the primary culture was transferred into 1 L of terrific broth (TB) and grown under identical conditions until the $OD_{600}$ reached ~0.8. Protein expression was induced with 1 mM isopropyl β-D-1-thiogalactopyranoside (IPTG) for 4 h at 37°C. Cells were harvested by centrifugation at 8,000 rpm for 30 min.

For protein purification, the bacterial pellet was resuspended in the lysis buffer containing 20 mM Tris-HCl, pH 8.0, with a protease inhibitor cocktail tablet (Roche) and subsequently lysed using a probe sonicator (Sonics & Materials Inc., pulse of 2s on, 1s off; 50% amplitude) for 15 min on ice. Following sonication, inclusion bodies were isolated from the cell lysate by centrifugation at 15,000 rpm for 45 minutes and were washed with 0.5% Triton X-100. The centrifugation, followed by the Triton X washing step, was repeated twice, and the pellet was dissolved in a solution of 8 M Urea containing 2% (v/v) β-mercaptoethanol (BME) to denature all the proteins present in the pellet. The resulting solution was dialysed using a 10 kDa molecular weight cut-off dialysis membrane against 20 mM Tris-HCl, at pH 8.0 and 4°C to slowly remove urea and β-mercapto-ethanol from the protein solution. The protein solution was then centrifuged at 15,000 rpm for 1h to pellet down any impurities. The supernatant of GH/PRL was loaded into the HiTrap anion exchange column, connected and operated through the FPLC system (GE Healthcare). The protein was eluted at the salt gradient of 0.5 M NaCl in 20 mM Tris-HCl pH 8.0 buffer (containing 0.01% sodium azide) and further purified by size exclusion chromatography (Superdex$^{TM}$ 200, GE) to isolate the monomeric fraction. For ΔPRL(108–116), the supernatant was passed through a 0.45 μm membrane filter and subsequently concentrated using Amicon centrifugal concentrators with 10 kDa and 50 kDa molecular weight cut-off membranes. The protein solution was further purified by size-exclusion chromatography using a Superdex™ 200 column. Fractions containing the monomeric protein were pooled and further concentrated using a centrifugal concentrator with a 10 kDa molecular weight cut-off. The purity of all purified proteins was checked using SDS-PAGE and MALDI-TOF. The concentration of the protein was measured using 280 nm UV absorbance, considering the molar absorptivity (ε) value of the PRL/ΔPRL(108-116), as 21805 $M^{-1}cm^{-1}$ and GH as 21170 $M^{-1}cm^{-1}$. The α-helical secondary structure of the purified protein was confirmed by CD. The monomeric protein fractions were immediately frozen in liquid nitrogen and lyophilised. The lyophilised protein powder was stored at -20°C until further use.

**Fluorescent labelling of proteins/peptides**

The NHS-Rhodamine and Fluorescein Isothiocyanate (FITC) labelling of proteins/peptides was performed as per the manufacturer's protocol (ThermoFisher Scientific, USA). Briefly, NHS-Rhodamine was dissolved in DMSO, was added as a 5-fold molar excess to the protein/peptide solution and was incubated in the dark on a magnetic stirrer with slow rotation for the first 1 hour at room temperature, followed by overnight incubation at 4°C. For FITC labelling, the protein/peptide solution was first buffer-exchanged into a pH 9, bicarbonate buffer, followed by performing the labelling reaction as per the manufacturer's instructions (ThermoFisher Scientific, USA). The dye-labelled protein solution was separated from the excess unreacted dye by using a PD10 column (GE Healthcare Life Sciences). The concentration of the dye-labelled protein and the degree of labelling were calculated as per the manufacturer's protocol, considering the molar extinction coefficients of NHS Rhodamine and FITC are 80,000 $M^{-1}$ $cm^{-1}$ and 70,000 $M^{-1}$ $cm^{-1}$, respectively.

**In-vitro phase separation assay**

For investigating the in-vitro phase separation, all protein/peptide solutions were diluted to a final protein/peptide concentration of 2 mg/mL in 20 mM phosphate buffer (PB), containing 0.01% sodium azide, and the pH was adjusted to either pH 7.4 or pH 6, based on experimental conditions. For GH and PRL, phase diagrams were constructed across a range of protein concentrations under varying conditions, including the presence of GAGs, $Zn^{2+}$, and different pH levels. The stock of 5 mg/mL GAGs (Heparin/CSA/CSB) and 40% stock PEG-8000 solution were prepared in 20 mM PB buffer (containing 0.01% sodium azide), pH 6. A 5 mM stock solution of $ZnSO_4$ was prepared in 20 mM Tris buffer (containing 0.01% sodium azide), pH 6, to avoid its precipitation in PB buffer. For GH and PRL phase separation, the phase regimes were constructed at various protein concentrations with changing additives (GAGs/$Zn^{2+}$/pH). For the in-vitro phase separation assay, protein/peptide solution was mixed with the respective NHS-rhodamine-labelled proteins/peptides at a molar ratio of 1:100 of labelled to unlabelled proteins/peptides and respective additives were added to induce phase separation inside test tubes. 50 μL of the phase-separated solution was further incubated in a small glass-bottom chamber, sealed with parafilm and kept inside a moist chamber in a 37°C incubator. The solution was monitored over time using an Olympus FV3000 laser scanning confocal microscope (100×/1.4 NA oil immersion objective) in the DIC (Differential Interference contrast) mode and fluorescence mode using a 561-nm DPSS 561-10 laser.

**Fluorescence recovery after photobleaching (FRAP)**

Phase-separated condensates of protein/peptides were subjected to time-dependent FRAP experiments, where all protein condensates were comprised of 1% molar ratio of NHS-rhodamine-labelled protein to the unlabelled protein. The FRAP experiment was performed on the Olympus FV3000 laser scanning confocal microscope, where a 50 μL phase-separated solution incubated inside a glass-bottom chamber was placed on a 100×/1.4 NA oil immersion objective.

A region of interest (ROI) within a condensate was photobleached using a 561-nm DPSS 561-10 laser at 100% laser power, while two additional ROIs of identical diameter were selected for background and passive-bleaching correction. Fluorescence intensities for all ROIs were recorded using the in-built software, and time-lapse images were acquired at a frame size of 512 × 512 pixels with 8-bit depth. The fluorescence-recovery profile, after background correction and normalisation, was fitted to a single-exponential growth function following previously established protocols (11).

The following equation was used for fitting as per the established protocol

$$I\,(t) = A\,(1 - exp\,(-t/\tau)) + C$$

Where τ indicates the time constant of fluorescence recovery, 'A' corresponds to the mobile fraction of the fluorescent probe, and '$C$' is the Y-intercept of the recovery profile. The mobile fraction (M.F.) of the proteins in the ROI was calculated by;

$$M.F = \frac{I_{\infty} - I_C}{I_{C_0} - I_C}$$

$I\infty$ = end value of fluorescence intensity post-recovery

The graphs were generated using OriginPro 2021 software (Origin Lab, USA).

**Static light scattering measurements**

The static light scattering experiment was performed to monitor the kinetics of phase separation of GH and PRL condensates at pH 7.4 with CSA and at pH 6 without any additives. For both conditions, 500 μM protein was used with or without 500 μM CSA inside a fluorescence cuvette, and the light scattering measurement was initiated immediately after inducing phase separation under respective conditions. The measurements were carried out on a JASCO FP-8500 (USA) spectrofluorometer using both excitation and emission wavelengths set to 350 nm with a 5-nm slit width. The scattering signal was recorded in continuous mode at 1-minute intervals. Each experiment was repeated twice. The scattering profile was plotted using Origin Pro 8 software (Origin Lab, USA).

Similarly, the effect of pH on GH phase separation was investigated using static light scattering at a wavelength of 350 nm. For this purpose, an initial scattering value at 350 nm was recorded in a JASCO FP8500 (USA) spectrofluorometer for a 200 μL phase-separated solution of 500 μM GH at pH 6. Then, the pH of the solution was gradually increased by 0.5 units, and the respective scattering value was recorded. After reaching pH 7.4, the pH was again gradually decreased to pH 6, and the static light scattering value at 350 nm was measured at each pH change by 0.5 unit. The experiment was repeated twice.

**Dense phase and dilute phase concentration determination**

To measure the concentration of the dense and dilute phases of the GH and PRL phase separation, the 500 μL solution of GH/PRL (500 μM) was adjusted to pH 6 to induce phase separation and incubated for 1 hour. The solution was then centrifuged at 15000g for 30 minutes. The supernatant was isolated completely, and protein concentration was measured based on the 280 nm absorbance of the protein. To measure the dense phase concentration, a minimum volume of pH 8.5 Tris buffer was added, which completely dissolved the pellet. The concentration was then calculated using 280 nm absorbance, taking into consideration the volume added to dissolve the pellet. Three independent experiments have been performed.

**Condensate mixing experiment**

To investigate the liquid-to-solid transition property of GH/PRL condensates at pH 7.4 (with CSA) and at pH 6, the molecular exchange property of GH/PRL condensates was tested over time. For that, two equal-volume (50 µL) phase-separated solutions of the same protein, formed under identical conditions, but one labelled with NHS-Rhodamine and the other labelled with FITC (1% molar ratio to unlabelled protein), were mixed immediately after phase separation (0 h) and after 1h of incubation at 37 °C in a moist chamber. The final mixture (100 µL) was drop-casted in a glass-bottom dish (SPL Lifesciences, Korea) and visualised under confocal microscopy (Olympus Fluoview FV3000 laser scanning confocal microscope (inverted) equipped with an iPlan-Apochromat 100X/1.4 NA oil immersion objective) using 488 nm and 561 nm OBIS lasers, keeping the same settings for all image acquisition. The confocal images of the yellow condensate population suggested complete molecular mixing of differently labelled proteins within condensates, while individual red/ green condensate populations suggested viscoelastic transition/solidification. The experiment has been repeated twice with similar observations.

**Interaction of PRL with Galanin (GAL) and Ribonuclease A (RNase A)**To examine the interaction of PRL with the specific interacting hormone GAL (known to be co-stored with PRL (12)) and a constitutive secretory protein, RNaseA (13), 500 µM PRL was mixed with either GAL (500 µM) or RNaseA (500 µM). Phase separation of the mixed solutions was induced under two experimental conditions: (i) pH 6 and (ii) pH 6 with 500 µM heparin. For each condition, the respective individual proteins were also checked for their phase separation. For imaging, 50 µL of the phase-separated solution was incubated in a small glass-bottom chamber, sealed with parafilm and incubated inside a moist chamber in a 37°C incubator. Imaging was carried out on an Olympus FV3000 laser-scanning confocal microscope using a 100×/1.4 NA oil-immersion objective, with fluorescence excitation provided by a 561-nm DPSS 561-10 laser. The experiment was performed twice, yielding similar observations.

**Thioflavin S (ThS) Staining**

Phase-separated solutions of all ten peptides and GH/PRL were tested for the ThS dye binding assay under various phase separation conditions at different time points. For this, 5 µl of 0.0625% ThS (w/v), prepared in 20 mM sodium phosphate buffer (pH 7.4), was mixed with 50 µl protein solution under phase-separate conditions and incubated in a sealed glass-bottom chamber. The confocal microscopy imaging was performed using an Olympus FV3000 laser scanning confocal microscope equipped with a 100×/1.4 NA oil immersion objective with 488 nm and 561 nm OBIS lasers, keeping the same settings for all image acquisition at different time points. The images were analysed using ImageJ (NIH, Bethesda, USA) software. To quantify the fluorescence enhancement, the mean fluorescence intensity of images was evaluated and plotted according to time.

**Thioflavin T (ThT) aggregation kinetics**

The aggregation kinetics of GH and PRL in various phase-separating conditions were monitored through amyloid-specific ThT fluorescence enhancement using a NUNC 96-well optical bottom plate on VANTAstar Plate Reader (BMG LABTECH, Germany) at 37°C. All GH and PRL phase separation solutions of 200 µL were set for ThT aggregation in the plate reader in such a way that the final concentration of PRL and GH is 500 µM, and the final concentration of ThT is 20 µM in the phase separating solutions. The experiment was initiated immediately after preparing the reaction mixture by monitoring fluorescence at 482 nm at 5-minute intervals, under non-shaking conditions, until the saturation of ThT intensity was reached. Three independent experiments were performed for each sample. The acquired kinetics data were plotted using Origin (2021b).

**Transmission Electron Microscopy (TEM)**

The morphology of protein/peptide hormone condensates was observed through TEM at different timepoints. For sample preparation, 10 µL of the phase-separated solution was drop-cast on a clean parafilm, and a copper formvar EM grid (Electron Microscopy Sciences, USA) was placed on top of the solution for 5 minutes of incubation. The grids were subjected to staining with uranyl formate (1% w/v) for 1 minute. The excess liquid was removed by carefully soaking through Whatman filter paper, and the grids were dried in a desiccator to remove all the moisture. The prepared grids were dried under an IR lamp before imaging. TEM images were acquired using a 200-kV transmission electron microscope (JEOL JEM 2100F, Japan) at 10,000× magnification, and the micrographs were captured digitally using the Gatan Microscopy Suite (Gatan, USA)

**Fourier-transform infrared (FTIR) spectroscopy**

FTIR was performed to examine the secondary structure change of protein/peptide hormones from monomers to the phase-separated condensates. For sample preparation for monomers, 10 µl of monomeric protein/peptide solution was spotted on a KBr pellet. To elucidate the peptide secondary structure of the condensates, the dense phase was isolated from a 50 µl phase-separated solution through centrifugation and spotted on the KBr pellet without dilution. All KBR pellets were dried under the IR (infra-red) lamp, and FTIR spectra were acquired using the Vertex 80 FTIR instrument, equipped with a DTGS detector (Bruker, Leipzig, Germany), in the amide I stretching frequency range of 1800–1500 cm−1 using an average of 32 scans at a resolution of 4 $cm^{-1}$. The recorded spectrum, corresponding to the wavenumbers 1700−1600 cm−1, was baseline-corrected, deconvoluted using the Fourier self-deconvolution (FSD) method, and then fitted with a Lorentzian function at OPUS-65 software (Bruker, Germany) according to the manufacturer's instructions. (14). The experiment was repeated two times with similar observations.

**X-ray fibril diffraction**

X-ray diffraction was performed for GH (with $Zn^{2+}$, pH 6) and PRL (with CSA, pH 6) phase-separated solutions under conditions that showed the highest ThT binding and the amyloid-specific cross-β peak in FTIR spectra. For sample preparation, 100 µL of the phase-separated solution after 24 h of incubation was ultracentrifuged at 90,000 rpm for 1 hr to isolate the dense phase. The pellet containing GH aggregates was redissolved in 5 µL of pH 6 Tris buffer and loaded into a pre-dried, clean 0.7 mm capillary tube, which was then dried overnight under vacuum. The entire capillary, containing the dried protein film, was placed in the path of the X-ray beam at 1.2 kW for a 5-minute exposure. The resulting image was acquired using a Rigaku R Axis IV++ detector (Rigaku, Japan) mounted on a rotating anode. The distance between the sample to the detector was kept at 200 mm, and the resulting image files were analysed and processed using Adxv software (Scripps Research Institute, USA).

**Monomer release assay**

The monomer release assay was performed as per the established protocol for the hormone amyloid release assay (12, 15, 16). For the experiment, we used GH and PRL amyloid-containing condensates, formed under the conditions $Zn^{2+}$ (pH 6) and CSA (pH 5.5), respectively, that showed the highest ThT and ThS binding, the amyloid-specific cross-β peak in FTIR spectra and the amyloid-specific diffraction pattern. For the release assay, a 100 µl aliquot of the GH (500 µM) and PRL (500 µM) phase-separated solution was ultracentrifuged at 90,000 rpm for 1 hr to isolate the dense phase fraction containing solid-like condensates. The pellet was resuspended in 50 µL pH 6 buffer and transferred into a modified PCR tube with a pierced hole in its cap. The modified PCR tube was sealed with a 50 kDa molecular weight cutoff membrane (Pierce, USA) and was placed inside a 2 ml cryotube (Nunc, Denmark) containing 1 mL of pH 7.4, Tris-HCl buffer with 0.01% sodium azide. The whole setup was then placed at 4 °C to prevent evaporation, and at a suitable time interval, 100 μl solution was taken from the released medium in the cryotube to measure 280 nm absorbance. After the measurement, the solution was returned to the cryovial to avoid any volume difference. An identical experimental setup was also checked, through the absorbance, keeping the buffer both inside and outside of the membrane, to determine whether degradation of the dialysis membrane is interfering with the assay. 500 μM monomer was also kept in another similar setup as a positive control to check the monomer release outside of the membrane through 280 nm absorbance. Similarly, to compare with disease-related solid condensates containing amyloid fibrils, a 500 μM α-Synuclein (α-Syn) solidified condensate was also put in another identical release setup. The secondary structure of the aggregated protein and the released protein from condensates was also checked by CD spectroscopy. Two independent experiments were performed for each sample.

**Circular Dichroism spectroscopy (CD)**

For CD measurement, the protein/peptide solutions were diluted in a suitable buffer to a final volume of 200 μl, resulting in a final protein/peptide concentration of 10-20 μM. The experiments were performed using a JASCO 810 CD instrument, where the sample was loaded in a quartz cell with a 0.1 cm path length (Hellma, Forest Hills, NY). The spectra were collected in the far UV range of 195–250 nm wavelength at 25°C. The raw data were processed as per the manufacturer's instructions. Three independent experiments were performed for each sample.

**Cell proliferation assay using Nb2 cells**

Nb2 cells were maintained in plastic culture dishes in RPMI medium (HiMedia, India) supplemented with 10% heat-inactivated fetal bovine serum (Gibco, USA), 10% horse serum (HiMedia, India), and 1× antibiotic solution. Cultures were incubated at 37 °C in a humidified atmosphere containing 5% $CO_2$. For the proliferation assay, 50,000 cells per well were seeded into 96-well plates using RPMI medium supplemented with 10% fetal bovine serum and 10% horse serum, followed by a 24-h incubation. Cells were then treated with GH monomer, PRL monomer, or hormone released from the corresponding amyloid condensates at a final concentration of 50 μM. An unrelated protein, γ-Globulin, served as a negative control. After 72 h of incubation, cell proliferation was assessed using the MTT assay. For this, 20 μL of MTT solution (5 mg/mL in PBS) was added to each well, and the plate was incubated for 4 h at 37 °C to allow formazan crystal formation. Formazan crystals were solubilised by adding 100 µL of acidified 10% SDS solution per well, followed by overnight incubation at 37 °C. Absorbance was measured at 570 nm with a reference wavelength of 650 nm using an ELISA plate reader. Fold changes in cell proliferation relative to untreated controls were then calculated and plotted using GraphPad Prism 8 software. All experiments were performed in triplicate.

**Dot blot with in vitro condensates**

To determine the amyloid oligomers and fibrils formation during phase separation and subsequent liquid-to-solid transition, the phase-separation mediated GH and α-Syn aggregation pathway was monitored through time-dependent dot blot with amyloid fibril-specific OC antibody (17) and oligomeric epitope-binding A11 antibody (18). For the experiment, GH phase separation was induced under the conditions $Zn^{2+}$ (pH 6), which showed amyloid aggregation as characterised by other biophysical techniques. Similarly, α-Syn phase separation was induced in the presence of PEG-8000, which was shown to form amyloid in previous reports (11, 19). During the time of phase separation and subsequent liquid-to-solid transition (up to 12 h), a 2 µL sample solution was spotted on a nitrocellulose membrane (Immobilon-NC, Millipore) at different time intervals and allowed to air-dry for 10 minutes. Once all spotting is completed, the membrane was then washed twice (2 × 8 minutes) with PBST (137 mM NaCl, 2.7 mM KCl, 10 mM $Na_2HPO_4$, 2mM $KH_2PO_4$, and 0.1% Tween), followed by blocking with 5% non-fat milk (HiMedia, Mumbai, India) for 1 hour. The membrane was then incubated overnight at 4°C with the corresponding primary antibody (OC or A11 antibody, 1:500 dilution) and washed with PBST (2 × 5 minutes) the next day. The membrane was then incubated with the corresponding horseradish peroxidase-conjugated secondary antibody (1:2000 dilutions) for 2 h at room temperature. Three PBST washes were performed (3 × 5 minutes), and the blot was developed by using the Western ECL detection kit (catalogue no. 1705060, Bio-Rad) according to the manufacturer's instructions. The images were captured using the Image Quant LAS 500 (GE Life Sciences, USA). Two independent experiments were performed.

**Intermolecular interaction study**

To elucidate the nature of intermolecular interactions underlying GH/PRL phase separation, various titration experiments were performed on preformed GH/PRL condensates as per the previously established protocol (20). To do this, preformed condensates at pH 6 were treated with either increasing concentration of NaCl (0-200 mM) or 1,6-Hexanediol (0-15% w/v). This is done to disrupt electrostatic and hydrophobic interactions, respectively. After each addition, the static light scattering reading at 350 nm was noted using a Spectrofluorometer (JASCO FP-8500, Japan) and plotted using GraphPad Prism 8 software. Further, the condensates, labelled with 1% NHS-rhodamine-conjugated protein, were also treated similarly and imaged using an Olympus FV3000 laser scanning confocal microscope (100×/1.4 NA oil immersion objective), using a 561-nm DPSS 561-10 laser. The experiment has been repeated twice with similar observations.

**Cell lines and transfection studies**

Mouse neuroendocrine cell line AtT-20 was kindly gifted by Marilyn Perrin, Salk Institute. AtT-20 cells were maintained in Dulbecco's Modified Eagle's Medium (DMEM) supplemented with 10% heat-inactivated Fetal bovine serum (FBS, Gibco, USA), 15% horse serum (HS, HIMEDIA, USA) and 1X Pen-Strep antimicrobial solution (HiMedia, India) at 37°C in a 5% $CO_2$ humidified incubator. The NIH3T3 mouse embryonic fibroblast cell line, a non-endocrine cell line, was maintained in DMEM, supplemented with 10% heat-inactivated FBS and treated with 1X Pen-Strep antimicrobial solution at 37°C in a humidified incubator containing 5% $CO_2$. For all live cell experiments, cells were seeded in the 35 mm confocal dishes at 70% confluency. For all immunostaining experiments, cells were seeded onto 18-mm glass coverslips in a 12-well plate at 70% confluency.

The transient transfection on the 70% confluent AtT-20 cells was performed by adding 1 μg of various plasmid constructs; namely GH-EGFP (Full-length GH with signalling sequence at N terminus of EGFP in pEGFP-N1 vector), PRL-EGFP (Full-length PRL with signalling sequence at N terminus of EGFP in pEGFP-N1 vector), pEGFP-N1 vector plasmid, ΔGH1 (deletion of 7-19 a.a), ΔGH2 (deletion of 99-113 a.a.), ΔPRL1 (deletion of 10-23 a.a.), and ΔPRL2 (deletion of 108-116 a.a.), ΔΔPRL (deletion of 10-23, 108-116 a.a.) plasmid; using Lipofectamine and P3000 following the manufacturer's protocol (L3000-008, Invitrogen, USA). Transient transfection with the GH-EGFP plasmid was also performed in NIH3T3 cells, similarly using Lipofectamine and P3000.

**Time-lapse confocal microscopy imaging**

Live cell imaging of transfected cells was performed at 16, 24, and 48 h post-transfection using the Olympus FV3000 laser scanning confocal microscope (100×/1.4 NA oil immersion objective) at a frame size of 1024 x 1024 pixels with 8-bit depth. The cell media was changed into 1 mL of Opti-MEM (catalogue no. 11058021, Gibco) before imaging, and 2 µL of Hoechst nuclear stain (catalogue no. 33342, Enzo, USA) was added to it. For all live-cell imaging, cells were incubated at 5% $CO_2$ and 37°C during the imaging. The excitation source of a 488-nm solid-state laser was used to monitor EGFP emission, 405 nm solid-state laser was used for Hoechst emission. All Z-stack images and movies were processed using ImageJ (NIH, Bethesda, USA) software.

The time-dependent (16 h, 24 h and 48 h post-transfection) ferret diameter of individual GH/PRL condensates was calculated using FIJI software from the nuclear plane confocal images (n=10 at each condition) of transfected AtT-20 cells within the thresholding range 0.4 µm - 3 µm (the lowest threshold was set above the confocal diffraction limit, and the highest threshold was set to remove the large condensate clusters). The condensate diameter distribution at each time point was plotted using GraphPad software.

**Lattice light-sheet microscopy**

Lattice light-sheet microscopy (LLSM) was performed to visualise the highly dynamic GH-EGFP condensates in the transfected AtT20 cells. For the experiment, the cells were seeded on 5-mm coverslips to reach ~70% confluency before transfection. 24-hour post-transfected cells were visualised under the 3i Lattice light-sheet microscope (3i, Colorado, USA), following the previous report (21). Briefly, a square lattice pattern was generated for the 488-nm channel using 51 beams spaced 0.99 µm apart (cropping factor 0.150), and volumetric stacks were acquired in dithered lattice mode with the same cropping factor. Samples were scanned in 0.3-µm steps, producing a deskewed step size of 0.163 µm. Excitation power at the image plane ranged from 0.14 to 0.43 mW to maintain an optimal signal-to-noise ratio. Emission was collected using a custom 488–640T/560R dichroic and a quad-notch filter (Semrock FF01-446/523/600/677), and images were recorded on a Hamamatsu ORCA-Fusion BT sCMOS camera with a 10-ms exposure time. Image volumes typically covered ~50 × 20 × 25 µm (X–Y–Z), corresponding to 140–200 planes, and were captured within ~2.5 s. Raw datasets were cropped and then deskewed in SlideBook to correct for the ~32° acquisition angle inherent to LLSM geometry. Deconvolution was performed using the Richardson–Lucy constrained iterative algorithm (10 iterations) with a theoretical PSF and a Gaussian smoothing radius of 0.3. Final images were prepared for display using FIJI.

**Immunofluorescence Staining**

30 hours post-transfection, cells were washed twice with phosphate-buffered saline (PBS, pH 7.4) and fixed with ice-cold 4% paraformaldehyde (PFA) in PBS for 15 minutes at room temperature in the dark. After fixation, permeabilisation was performed using 0.2% Triton X-100 in PBS for 5 minutes at room temperature, followed by incubation with 5% bovine serum albumin (BSA) in PBS for 1 hour at room temperature to block non-specific binding. The coverslips were then incubated with various primary antibodies, namely mouse monoclonal anti-Rabbit CgA (Abcam, ab254322) antibody (1:250), rabbit monoclonal anti-TgN38 (Abcam, ab283678) antibody (1:500), mouse monoclonal anti-ACTH (Invitrogen, MA5-38437) antibody (1: 250), and rabbit polyclonal anti-β endorphin (Abcam, ab5028) antibody (1:500), diluted in PBST (PBS with 0.1% Tween-20) containing 5% BSA and incubated overnight at 4°C in a humidified chamber. Following primary antibody incubation, cells were washed three times with PBST and incubated with Alexa Fluor 555, conjugated with either anti-mouse (Invitrogen, A11001) or anti-rabbit (Invitrogen, A21428) secondary antibody (1:500 dilution in PBST + 5% BSA) for 1 hour at room temperature on a rocker in the dark. Coverslips were then washed with PBST, and nuclei were stained with DAPI (1:1000 dilution from a 1 mg/ml stock) for 2 minutes at room temperature in the dark, followed by a final PBS wash. Coverslips were mounted onto slides using an antifade mounting medium (90% glycerol, 10 mM Tris pH 8.0, and 2.5% DABCO) and stored at 4°C. Imaging was performed using an Olympus FV3000 laser scanning confocal microscope (100×/1.4 NA oil immersion objective). The images were analysed using ImageJ (NIH, Bethesda, USA) software. Excitation/emission wavelengths were set at 488 nm/530 nm for GFP, 555 nm/565 nm for Alexa Fluor 555, and 358 nm/461 nm for DAPI. Images were processed and analysed using Fiji software (NIH). All immunostaining experiment has been performed twice with similar observation.

**Immunohistochemistry**

Adult Sprague–Dawley rats (200–250 g, both sexes) were housed under standard conditions, including a 12 h light/12 h dark cycle, with free access to chow and water. All procedures were approved by the Institutional Animal Ethical Committee (IAEC) of NISER, Bhubaneswar (protocol numbers NISER/SBS/AH-210 and NISER/SBS/AH-212) and were conducted in accordance with the Committee for Control and Supervision of Experiments for Animals (CPCSEA) guidelines. The sectioned pituitary gland, mounted on poly-L-lysine (Sigma) coated glass slides, was a kind gift from Prof. Praful Singru (National Institute of Science Education and Research). The sections were processed for double immunofluorescence and subjected to a xylene-alcohol gradient rehydration process to remove the embedding medium and restore hydration. The gradient involved sequentially immersing the sections in increasing concentrations of ethanol, followed by xylene and vice veThe sectioned pituitary gland, mounted on poly-L-lysine (Sigma) coated glass slides, was a kind gift from Prof. Praful Singru (National Institute of Science Education and Research).rsa, starting from 100% xylene, transitioning to decreasing ethanol concentrations (100%, 95%, 70%, and 50%), and finally to distilled water. Tissue permeabilisation was done by treating the sections with 0.25% Triton X-100 for 10 minutes. Following that, the sections were rinsed twice in TBST (TBS with 0.2% Tween-20) for 5 min. The sections were incubated in a blocking solution (5% BSA in TBST) for 1 hour. The double immunofluorescence was performed in a humidified chamber by incubating the sections in a mixture of two primary antibodies of ACTH (Invitrogen, MA5-38437, 1:250), GH (R&D Systems, dilution 1:100), and CgA (Abcam, ab254322, 1:250 in 5% BSA) diluted in the blocking solution overnight at 4 °C. The next day, sections were rinsed three times in TBST and further incubated with the secondary antibody of goat anti-mouse Alexa-488 (1:400), goat anti-rabbit Alexa-488 (1:500), or goat anti-mouse Alexa Fluor-555 (1:500) (Thermo Scientific, USA) for 2h at room temperature in a humidified chamber. The sections were washed with 50% ethanol for 2 min, followed by TBST washing for 3 min. The tissue sections were stained with 0.5% DAPI (Sigma-Aldrich) for 5 minutes in the dark, followed by mounting in a medium containing 90% glycerol, 10% PBS, and 1% DABCO (1,4-diazabicyclo[2.2.2]octane; Sigma-Aldrich). Images were acquired using an Olympus Fluoview FV3000 laser scanning confocal microscope.

**Chemical treatment in AtT-20 cells and modulation of hormone condensates**

AtT-20 cells cultured on confocal dishes were imaged 30 hours post-transfection using a confocal microscope (Olympus Fluoview FV3000 laser scanning confocal microscope) before any chemical treatment. To examine the effect of pH on GH condensates, cells were incubated for 2 hours with 200 μM chloroquine (C6628, Sigma Aldrich) dissolved in PBS, following a previously reported protocol (22), and subsequently imaged under identical acquisition settings as untreated controls. The intensity quantification of the soluble fraction (diffused EGFP signal across the cell) relative to the total mean cellular fluorescence was performed before and after chloroquine treatment using FIJI software. Briefly, for each 8-bit image, the total mean cellular intensity was measured by applying a threshold that encompassed the entire cell. Next, a second threshold was applied to selectively identify all puncta, and the resulting puncta ROIs were subtracted from the whole-cell ROI. The mean fluorescence intensity measured from this puncta-excluded region represented the soluble fraction. The ratio of soluble to total mean intensity was calculated for each cell (n = 7 per condition) and compared using the bar plot, plotted by GraphPad.

Further, to investigate the active transport of GH granules in AtT-20 cells, microtubule destabilisation was carried out by incubating transfected cells with 5 µM Nocodazole (dissolved in DMSO) for 2 hours and imaged under the confocal microscope using the same acquisition setting as untreated cells. The destabilisation of the microtubule resulted in random condensates clumping throughout the cells. The amyloid status of the GH granules inside AtT-20 was checked using Amytracker 630 dye (Ebba Biotech). To do so, at 30 hours post-transfection, the cell media was replaced with PBS containing Amytracker 630 (Ebba Biotech) (1:250 dilution), followed by incubation at 37 °C for 1 h. Cells were then washed twice with fresh PBS and incubated with culture medium containing Hoechst dye for nuclear staining. Fluorescence imaging was performed using the confocal microscope equipped with 488 nm and 561 nm lasers to detect EGFP and Amytracker signals, respectively. To assess the role of hydrophobic interactions in GH condensate stability, transfected AtT-20 cells were treated with 1.5% (w/v) 1,6-Hexanediol (added to the culture medium). After 30 minutes of incubation, cells were fixed with 4% paraformaldehyde (PFA) and imaged using the same confocal settings as untreated cells. All the experiments have been performed twice independently with similar observations.

**Subcellular soluble and insoluble fraction preparation**

To isolate soluble and insoluble protein fractions, $1*10^6$ AtT-20 cells were seeded in 35 mm culture dishes and transfected with the GH–EGFP plasmid at ~70% confluency using Lipofectamine and P3000, following the manufacturer's protocol. For the time-dependent experiment, cells were harvested at the indicated time points by trypsinisation, followed by washing with PBS twice. The cell pellet was resuspended in a fractionation buffer containing 0.32 M sucrose, 1.5% Lubrol, 0.5% BSA, 10 mM sodium phosphate (pH 7.0), and 1× protease inhibitor cocktail (PIC). The suspension was ultracentrifuged at 50,000 × g for 1 hour at 4 °C, yielding the Lubrol-soluble supernatant (S fraction) and Lubrol-insoluble pellet (P fraction). The pellet was subsequently resuspended in lysis buffer containing 62.5 mM Tris (pH 6.8), 2% SDS, and 10% glycerol. Both fractions were stored at –20 °C until further use for western blot analysis.

**Temperature-dependent study**

To examine the effect of temperature on intracellular GH phase separation and granule formation, 1 × $10^6$ AtT-20 cells were seeded in two confocal dishes (for live-cell imaging) and two 35 mm culture plates (for subcellular fractionation). Cells were transfected with the GH–EGFP plasmid using Lipofectamine and P3000, following the manufacturer's instructions. After transfection, one set of cells was incubated at 20 °C in a 5% $CO_2$ humidified incubator, while the control set was maintained at 37 °C under identical conditions. At 30 hours post-transfection, both test and control cells were imaged under a confocal microscope (Olympus Fluoview FV3000 laser scanning confocal microscope) using a 488 nm laser to visualise GH–f of TGN/non-TGN fluorescence intensity in these two temperatures was quantified using FIJI from 8-bit images (n=10, each condition) by applying respective threshold covering the TGN region to get TGN intensity, followed by subtracting that region from the whole cell to obtain non-TGN intensity. The ratio was plotted as a bar graph using GraphPad software. In parallel, subcellular fractionation was performed from both sets at the same time point using the previously described method, followed by Western blot analysis. The experiment has been repeated twice.

**Western blotting**

For Western blot analysis, 40 µg of protein (quantified using a NanoDrop spectrophotometer) from the fractionated cell lysates was separated by SDS–PAGE and transferred onto a 0.22-µm nitrocellulose membrane. After transfer, the membrane was blocked with 5% BSA in TBST (TBS containing 0.1% Tween-20) for 1 h at room temperature. It was then incubated overnight at 4 °C with an anti-GFP primary antibody (Invitrogen, catalogue No. 14-6674-82) diluted 1:1000 in 5% BSA/TBST. Following washes with TBST, the membrane was incubated for 1 h at room temperature with a goat anti-mouse IgG (H+L) HRP-conjugated secondary antibody (Merck, Cat. 401253) diluted 1:2000 in 5% BSA/TBST. After three additional TBST washes, signals were developed using the Clarity Western ECL kit (Bio-Rad, catalogue No. 1705060) according to the manufacturer's protocol. Images were acquired using an ImageQuant LAS 500 system (GE Life Sciences, USA) and quantified with Bio-Rad Image software. Each experimental condition was analysed using three independent Western blots.

**Immunoprecipitation and dot blot of the GH-EGFP-positive fraction of the cell lysate**

AtT-20 cells, post 30 h of transfection with the GH-EGFP plasmid, were harvested by trypsinisation. The cell pellets were washed twice with ice-cold PBS and lysed in an ice-cold lysis buffer containing 50 mM Tris, pH 7.4, 150 mM NaCl, 1 mM EDTA, 0.5% NP-40, and protease inhibitor cocktail. The cell pellets were vigorously resuspended in the buffer by pipetting and briefly vortexed before being kept on ice for 5-10 minutes. The cell suspension was then centrifuged at 13,000 rpm for 15 minutes at 4°C. Following this, the soluble supernatant fraction was transferred to a fresh microcentrifuge tube and maintained on ice until further processing. Meanwhile, protein G Sepharose beads were washed twice with ice-cold lysis buffer to equilibrate them and stored on ice as a 1:1 bead slurry. The cell lysates were then incubated with equilibrated beads for 1 hour at 4°C on a shaker/tube rotator. This was followed by brief centrifugation of the lysates at ~1000g for 5 minutes to pellet the beads, and the resulting "precleared" soluble supernatant was then transferred to a fresh microcentrifuge tube and incubated with the anti-GFP antibody (1:100) on a shaking platform overnight at 4°C. The next day, both reaction tubes were centrifuged at 1000 x for 5 minutes so as to pellet down the beads. The beads were then washed at least 5 times with ice-cold lysis buffer. After the final wash, the buffer was discarded, and the beads were incubated with 0.1 M Glycine (pH 4.0) at 60°C for 5 minutes. The tubes were again briefly centrifuged, and the supernatants were used for the dot blot assay as described before.

**Secretory granule isolation and immunoprecipitation of GH-EGFP condensates**

GH condensates were isolated from 40 hours post-transfected AtT20 cells, cultured in three 90 cm dishes following the reported protocol (15, 23) with slight modification. Briefly, the cells were washed twice with PBS buffer and pelleted down by centrifuging at 1000 rpm for 10 minutes. The pellet was resuspended in SG buffer (0.25 M sucrose in 20 mM Tris-HCl buffer, 1 mM EDTA, pH 7.2) and homogenised with 10 strokes using a glass-Teflon homogeniser (Sigma) on ice. The cellular homogenate was then centrifuged at 600 × g for 5 min, and the pellet containing intact cells and nuclei (P1) was washed with 25 ml of SG buffer followed by a second centrifugation at 600 × g for 5 min. The supernatant from both centrifugations (S1 and S2) was mixed and further centrifuged at 4000 x g for 10 min, which produced supernatant (S3) and pellet (P3). S3 was further centrifuged for 20 min at 22,000 × g, which produced the pellet (S4) containing cell mitochondria, lysosomes, lightweight granules, and dense granules. S4 was resuspended in 2 mL of SG buffer and washed twice again following the same procedure, reported to eliminate the possibility of cold-induced adhesiveness in the crude granule fraction. The subsequent steps were strictly performed at 4°C. 50% Percoll solution in SG buffer was centrifuged at 30,000 × g for 15 min at the Optima MAX-XP Ultracentrifuge (Beckman Coulter) in a 10 mL polycarbonate ultracentrifuge tube (Beckman Coulter) to prepare around 8 mL density gradient. Percoll is a biologically inert colloidal suspension of minute silica beads, which is high-density but low-viscosity in nature and is heavily used to isolate organelles (24). The final pellet solution, containing the crude granule fraction, was resuspended in 200 µL SG buffer and layered on top of the gradient, followed by centrifugation at 30,000 × g for 20 min. After centrifugation, a distinct white band appeared close to the top of the gradient, containing mitochondria, lysosomes, and lightweight granules (granule L). The upper band was removed by aspiration and further centrifuged at 100,000 × g for an additional hour, resulting in the formation of a lower band at the gradient bottom containing dense core granules (granule H). Each band was collected, washed twice with ice-cold SG buffer. The pellet was diluted two-fold with SG buffer and centrifuged at 100,000 × g for one hour. This high-speed centrifugation step significantly removed the Percol content from the granule fractions. The purified granules were further immunoprecipitated using the EGFP primary antibody following the previously mentioned protocol. To detach the bead from the immunoprecipitated granule fraction, the solution was incubated at pH 3.5, 25 mM Glycine buffer for 30 minutes at 56 °C, followed by centrifuging at 1000 rpm for 5 minutes. The isolated granule fraction was observed under the confocal microscope using a 488 nm laser to detect EGFP fluorescence. The tryptophan fluorescence and EGFP fluorescence were measured using a Horiba Fluoromax 4 fluorometer by exciting at 280 nm and 488 nm, respectively. The TEM grid was also prepared from the EGFP-enriched granule fraction as per the previously mentioned in previous section (Transmission Electron Microscopy), and images were acquired using a 200-kV transmission electron microscope (JEOL JEM 2100F, Japan) with 10,000× magnification.

**Immunoelectron microscopy of AtT-20 cell cross-section**

To visualise the morphology of GH condensates/secretory granules inside AtT-20, we performed immunoelectron microscopy on the ultrathin (50-70 nm) cross-sections of GH-EGFP-transfected AtT-20 cells. For that, 40 h post-transfection, cells cultured in two 90 mm dishes were washed twice with PBS and fixed with 3% glutaraldehyde (pH 7.4) for 4–6 h at 4 °C. Cells were then washed with 0.1 M cacodylate buffer twice and post-fixed in freshly prepared 1% (w/v) Osmium Tetroxide (OsO4) solution in cacodylate buffer (1:1) for 2 hrs at 4°C, followed by two washes with distilled water for 10 minutes each. The samples were dehydrated through a graded ethanol–acetone series (30%, 50%, 70%, 90%, and 100%) for 15–30 min at each step, followed by two washes with propylene oxide (PO) (10 min each). The cell pellet was treated serially with PO/Epon (1:1) and PO/Epon (1:3) for 30 minutes and finally in Epon (Epoxy resin/Araldite) for 1 hour. The sample was then placed in block moulds, embedded in Epon (Epoxy resin/Araldite), and heated in the oven at 60-65°C for 24-48 hrs to polymerise Epon. Prepared resin blocks were cut into ultrathin sections (50-70 nm thickness) using a diamond knife in the Leica Ultramicrotome system (EM UC7) and mounted on Formvar-coated copper grids for immunogold labelling.

Immunolabeling on thin cell sections, mounted on a grid, was performed following the previously reported protocol (25). Briefly, the section was blocked using 5% FBS solution in 10 mM glycine, pH 7.4 buffer for one hour, followed by anti-mouse EGFP primary antibody (Invitrogen, 14-6674-82) treatment in a 1:2000 dilution in blocking buffer containing 0.01% Tween 20 for overnight incubation at 4°C. The sections were washed with 10 mM glycine buffer three times and incubated with anti-mouse IgG antibody conjugated with 10 nm Gold particle (Sigma (G7777) with a 1:500 dilution for 2 hours at room temperature. The grid was washed three times with glycine buffer, and contrast was generated by staining with 5% (w/v) lead acetate for two minutes and 1% (w/v) uranyl formate for 2 minutes. The grids were finally washed in Milli Q water and kept under vacuum for complete drying. The grids were then observed under the 300 kV Themis 300 G3 transmission electron microscope (Thermo Scientific) for image acquisition.

**Fluorescence lifetime experiment**

The material properties of the intracellular nascent GH condensates, which are populated at the TGN, and the mature SGs, populated at the cellular tip, were studied by monitoring the EGFP fluorescence lifetime at these two populations. To investigate that, AtT-20 cells, seeded in 18 mm coverslips in the 12-well plate, were transfected with the GH-EGFP plasmid at 70% confluency of the cells. 30 hours post-transfection, cells were fixed using ice-cold 4% paraformaldehyde and a coverslip was mounted on a slide. Fluorescence lifetime imaging (FLIM) was performed to record the EGFP fluorescence lifetime from the immediately fixed cells using the MicroTime 200 (MT200) time-resolved confocal microscope using the time-tagged time-resolved (TTTR) methodology. The setup was attached to an inverted microscope (Olympus IX71) equipped with a water immersion objective (UPlanSApo NA 1.2, 60×, WD = 0.28 mm). The sample was excited using a 440 nm pulsed diode laser, and the emitted fluorescence passed through a 440/532 dichroic mirror, a 465 long pass filter, and a 50 μm pinhole to get rid of out-of-focus light. The collected fluorescence signal was directed to a single-photon avalanche photodiode (SPAD) detector. The ROI containing the TGN or Tip population was imaged at 512 × 512 pixel resolution with a collection time of 0.60 ms per pixel, using the in-built SymPhoTime 64 software (PicoQuant, Germany). The acquired lifetime decay was analysed using the multi-exponential reconvolution method in SymPhoTime 64 software (PicoQuant, Germany), taking into account the instrument response function (IRF) and background decay. The decay profiles were satisfactorily fitted to the biexponential equation:

$y(t) = A_1 \exp(t_1/\tau_1) + A_2 \exp(t_2/\tau_2)$

where, $A_{sum} = A_1 + A_2$

$\tau_{av\ amp} = (A_1\tau_1 + A_2\tau_2)/A_{sum}$

A = exponential prefactors (amplitudes)

$\tau$ = exponential decay time (lifetimes)

$A_{sum}$ = Fluorescence intensity at time zero

$\tau_{av\ amp}$ = Amplitude weighted average lifetime (mean decay time at time zero)

**Single-particle tracking**

To track the dynamics of GH condensates at the trans-Golgi network (TGN) and mature secretory granules at the cellular tip, single-particle tracking was performed at individual condensate/granule resolution. For that, single particle tracking information of the individual puncta from the TGN and the Tip was extracted from the time-lapse video of GH-EGFP transfected AtT-20 cells, post 30 h of transfection, using the TrackMate plugin in ImageJ software, where the Gaussian low-pass filter was applied to reduce the noise and improve the particle detection accuracy. The quality threshold setting parameter was adjusted manually for individual images. A simple LAP tracker was used, keeping the gap tolerance, a maximum of 2 frames. The extracted trajectories were further analysed using the msdanalyzer package (26) in MATLAB. We computed the Diffusion coefficient (D), the mean squared displacement (MSD), and the correlation of track straightness to diffusion exponent ($\alpha$) for individual particles as well as the average values for all particles from a single cell. The computed results of TGN and Tip population were plotted using Origin and GraphPad plotting software.

**$BaCl_2$-stimulated release experiment**

Mature SGs, stored at the cellular tip, undergo regulated exocytosis upon secretagogue stimulation, resulting in the burst release of hormones (27). To examine whether GH phase separation leads to the formation of functional SGs capable of regulated secretion, we used Barium Chloride ($BaCl_2$), a well-established secretagogue that induces hormone release by elevating intracellular calcium in AtT-20 cells (28). For this, $1\times10^6$ AtT-20 cells were seeded in two 35 mm culture dishes (for Western blot) and one confocal dish (for live-cell imaging) and transfected with GH–EGFP at ~70% confluency. After 40 h of transfection, cells were rinsed with PBS and imaged in Opti-MEM using an Olympus Fluoview FV3000 laser scanning confocal microscope with 488 nm laser excitation to record transfected cell images before the treatment. The medium was then replaced with Opti-MEM containing 1 mM $BaCl_2$, and cells were incubated for 30 minutes before imaging under the identical acquisition settings. The ratio of TGN to non-TGN EGFP fluorescence intensity before and after $BaCl_2$ treatment was quantified from 8-bit images (n = 10 cells per condition) using FIJI, as described above.

To assess dose-dependent release through western blotting, cells were rinsed with PBS three times after 40 h of transfection, followed by incubation in 200 µL Opti-MEM for 30 minutes to collect basal secretion. The media were then replaced with Opti-MEM containing 1 mM or 5 mM $BaCl_2$ (in two different setups), and samples were collected after 30 minutes of incubation. In all collected media, 1X protease inhibitor cocktail was added, followed by centrifuging at 15000 rpm for 1 hour to remove cell debris, and the samples were stored at -20 °C for further use. 20 µL of each solution was loaded in SDS-PAGE electrophoresis to process for Western blot using anti-GFP (Invitrogen, catalogue no: 14-6674-82) primary antibody. The fold change in GH-EGFP secretion in the presence of 1 mM and 5 mM $BaCl_2$ relative to the basal secretion was calculated from three western blots and plotted as a bar graph. Additionally, 100 µL of each sample was analysed using a Horiba Fluoromax-4 fluorometer (excitation at 488 nm; emission 500–550 nm) to quantify EGFP fluorescence intensity at 513 nm, which was plotted as a bar graph.

#### Supplementary Figures

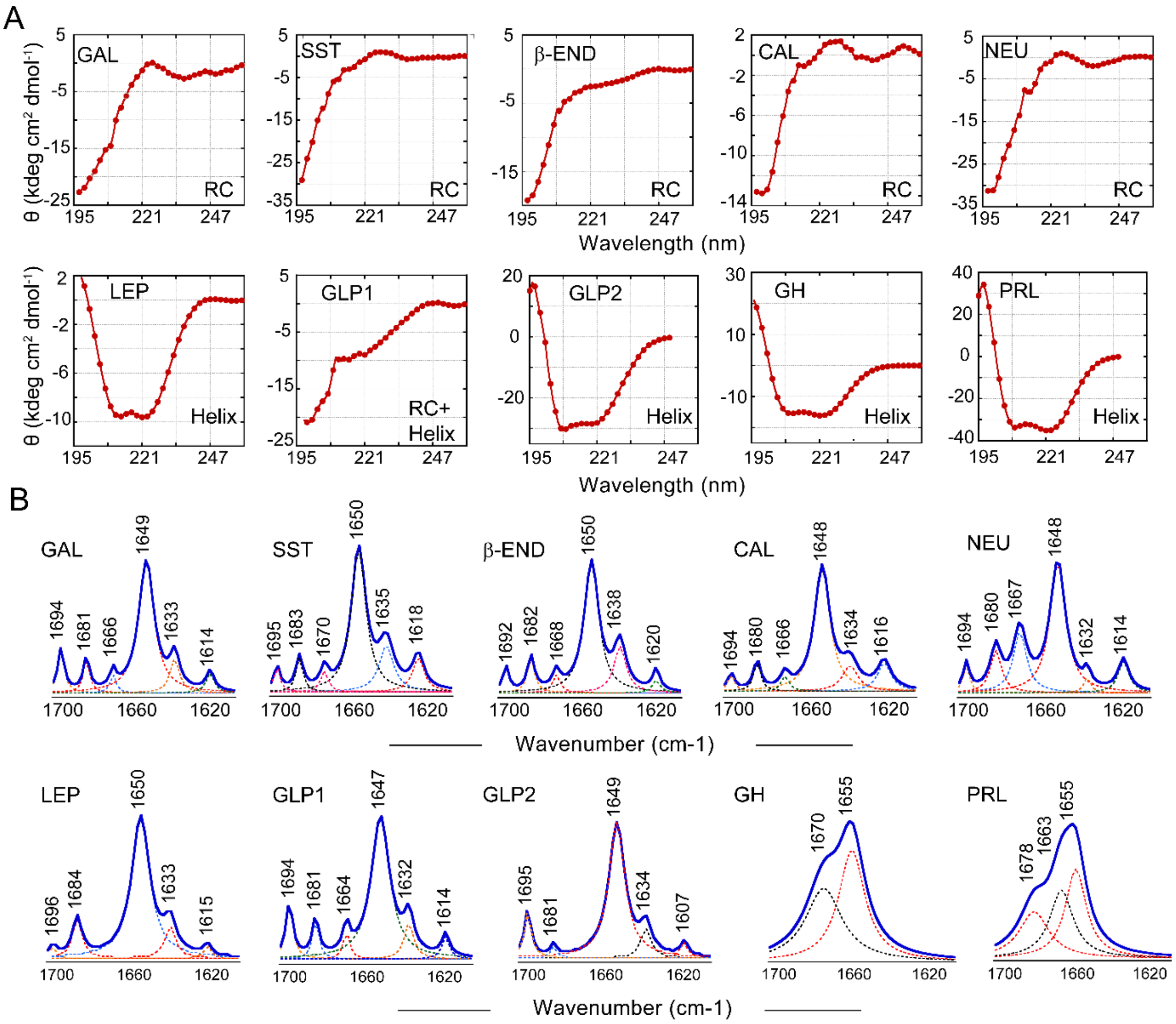


**Figure S1: Diverse secondary structure of protein/peptide hormones: (A)** CD spectroscopic analysis showing various secondary structures of protein/peptide hormones under study, immediately after dissolving in phosphate buffer (pH 7.4). While GAL, SST, β-END, CAL, NEU showed random coil (RC), LEP, GLP2, GH and PRL showed α-helix conformation, and GLP1 showed a mixture of RC and helix conformation. **(B)** Representative curve-fitted FTIR spectra (blue line) along with Fourier deconvoluted individual peaks (dotted lines), representing different secondary structures of protein/peptide hormones, immediately after dissolving in phosphate buffer (pH 7.4).

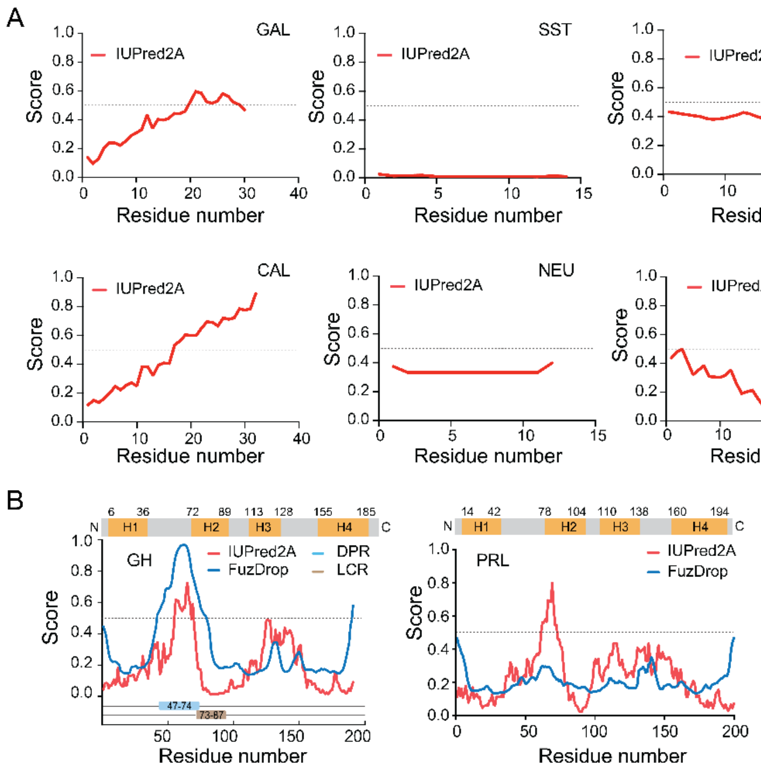


**Figure S2: In silico analysis of protein/peptide hormones: (A)** In silico analysis of peptide hormones using IuPred2A, predicting the intrinsic disordered region in the amino acid sequence. FuzDrop analysis for phase separation propensity prediction was performed only on Leptin (LEP), as it qualified for the amino acid length requirement. **(B)** Cartoon representation showing the four α-helix positions (H1-H4) in the GH and PRL amino acid sequence (top). In-silico analysis on GH and PRL using IuPred2A, FuzDrop and SMART showing the intrinsic disordered region, Droplet-promoting region (DPR) and low complexity region (LCR), respectively (bottom).

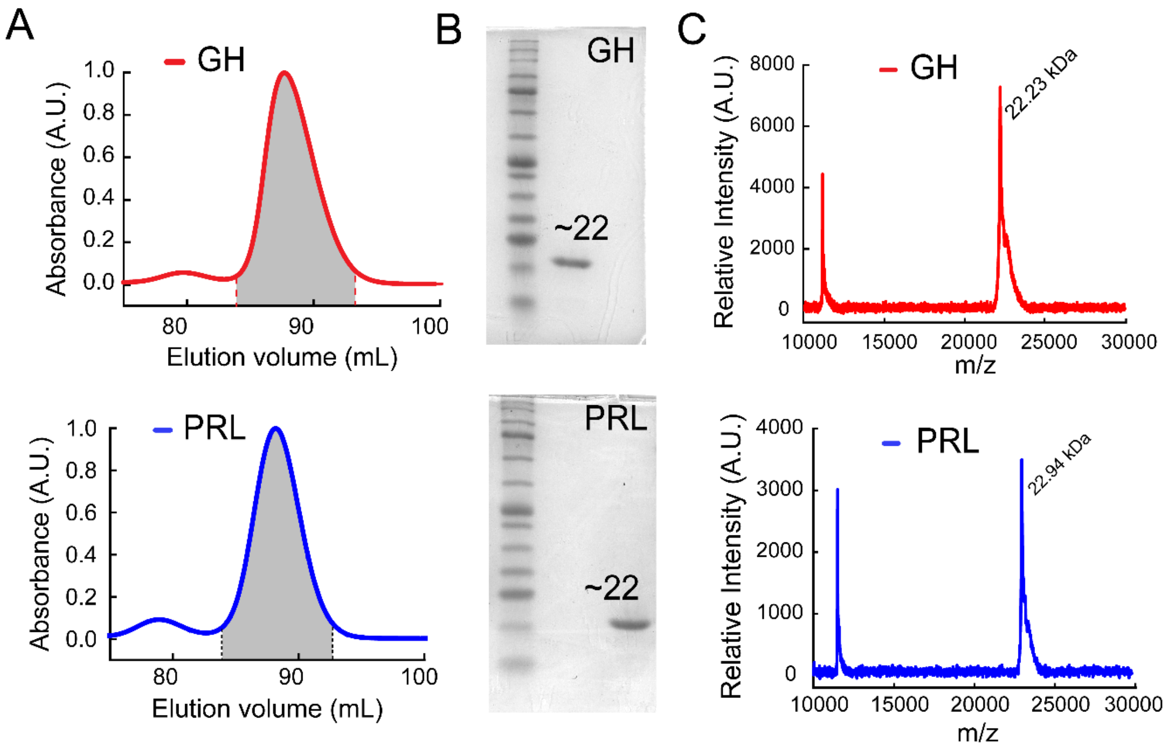


**Figure S3: Biophysical characterisation of expressed and purified GH and PRL. (A)** Size exclusion chromatogram of purified GH (top), PRL (bottom) after recombinant expression in BL21 (DE3) cells, and purification via anion exchange column chromatography. **(B)** The size exclusion proteins showed a single band at ~22 kDa in 15% SDS-PAGE, confirming the pure protein fraction of GH (Top) and PRL (bottom). **(C)** MALDI profile of GH (top) and PRL (bottom), confirming their molecular weight peak at m/z= 22.23 kDa and 22.94 kDa, respectively.

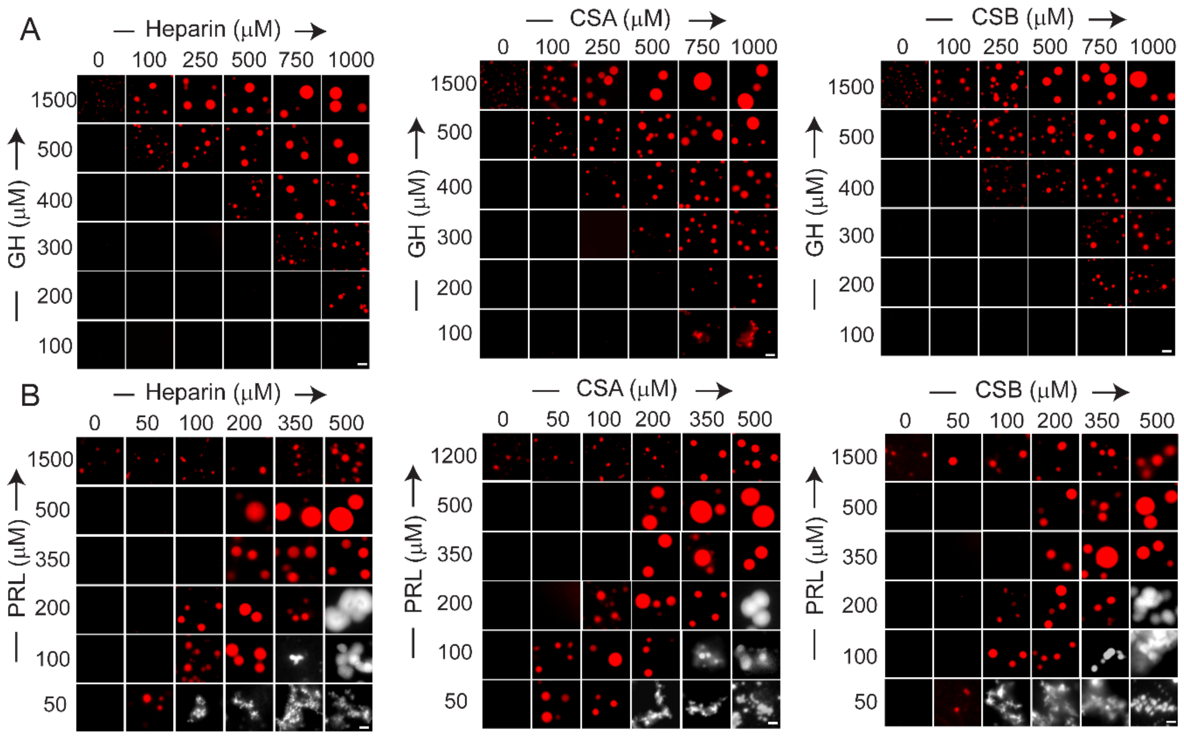


**Figure S4: Phase regime of GH and PRL at pH 7.4 at varying protein and glycosaminoglycan (GAG) concentrations. (A)** Phase regime of GH in the presence of CSA, CSB, and Heparin at pH 7.4, showing an increase in GAG concentration promoting GH Phase separation. **(B)** Phase regime of PRL in the presence of different GAGs: CSA, CSB, and Heparin at pH 7.4, exhibiting the highest phase separation propensity of PRL at a 1:1 ratio (protein: GAG), while a high GAG ratio promotes condensate clumping (grey scale). The scale representing 2 μm.

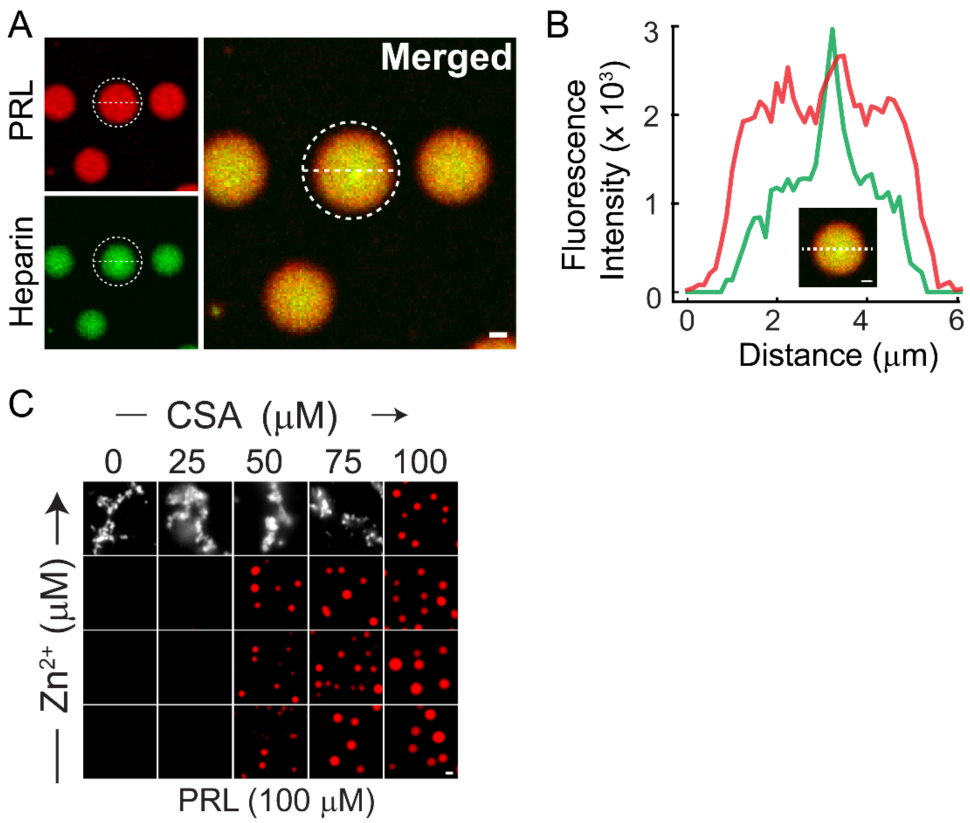


**Figure S5: Interaction of GAGs and $Zn^{2+}$ with GH/PRL in condensate. (A)** Fluorescence image showing distribution of labelled heparin (green) inside PRL (red) condensate. Scale bar: 2 µm. **(B)** Intensity profile plot of Heparin (green) and PRL (red), showing the distribution of both components inside a single PRL condensate. Scale bar: 1 µm. **(C)** Phase regime demonstrating the effect of $Zn^{2+}$ and PRL with 100 µM PRL (L), where the proteins otherwise do not form visible condensates. Scale bar: 1 µm.

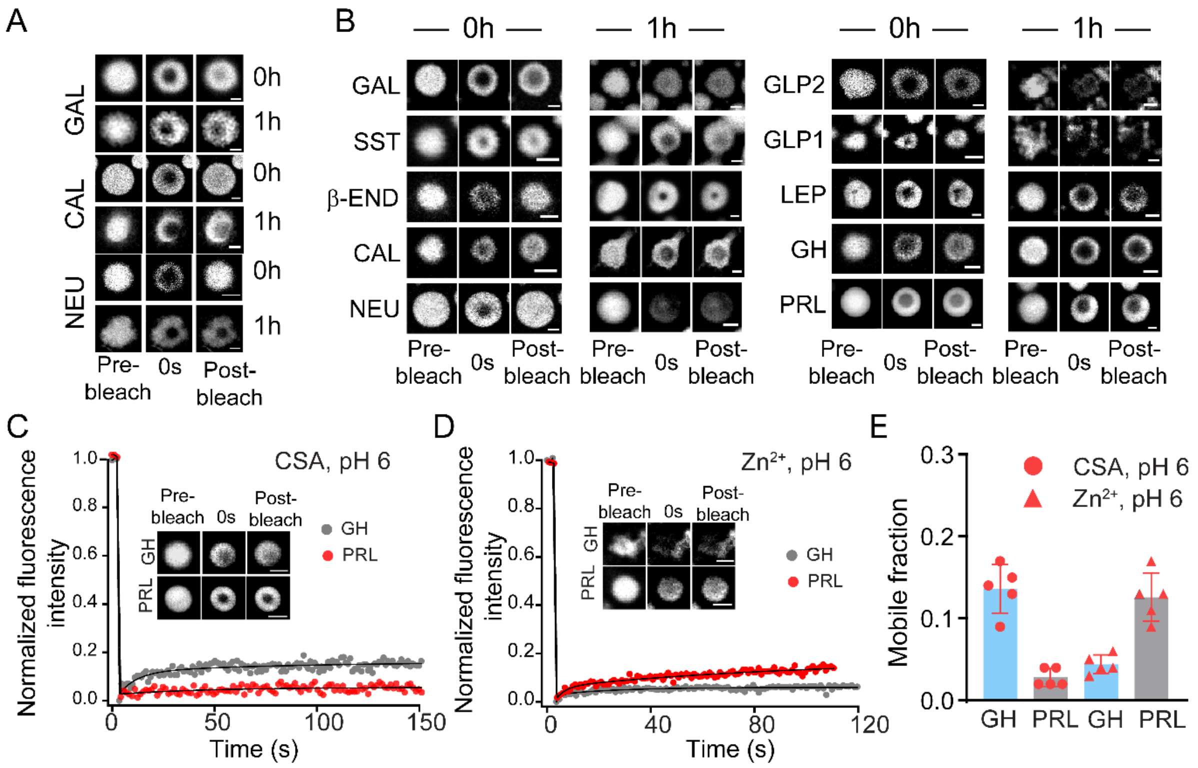


**Figure S6: Liquid-to-solid transition of hormone condensates: (A)** Representative grayscale confocal images showing FRAP at pH 6 peptide condensates (GAL, CAL, and NEU) immediately after formation (0 h) and after 1 h of incubation. For each condition, images of a single condensate are shown before photobleaching (Pre-bleach), at the time of bleaching (0 s), and during recovery up to 150 s (Post-bleach). A progressive reduction in FRAP recovery is observed over time. **(B)** Representative grayscale confocal images of peptide condensates formed at pH 6 in the presence of Heparin, imaged immediately after formation (0 h) and after 1 h of incubation during the FRAP experiment. Pre-bleach, 0 s, and Post-bleach (up to 150 s) images of single condensates demonstrate consistently low FRAP recovery even at 0 h. **(C, D)** Normalised FRAP intensity profiles of GH (grey) and PRL (red) condensates immediately after formation at pH 6 in the presence of CSA (**C**) or $Zn^{2+}$ (**D**), showing negligible fluorescence recovery following photobleaching. Representative grayscale images of single condensates before bleaching (Pre-bleach), at bleaching (0 s), and during recovery (Post-bleach) are shown, with a scale bar of 1 µm. **(E)** Mobile fraction calculated from five independent FRAP profiles for immediately formed GH (blue bars) and PRL (grey bars) condensates at pH 6, in the presence of CSA (circles) or $Zn^{2+}$ (triangles). All scale bar: 1 µm.

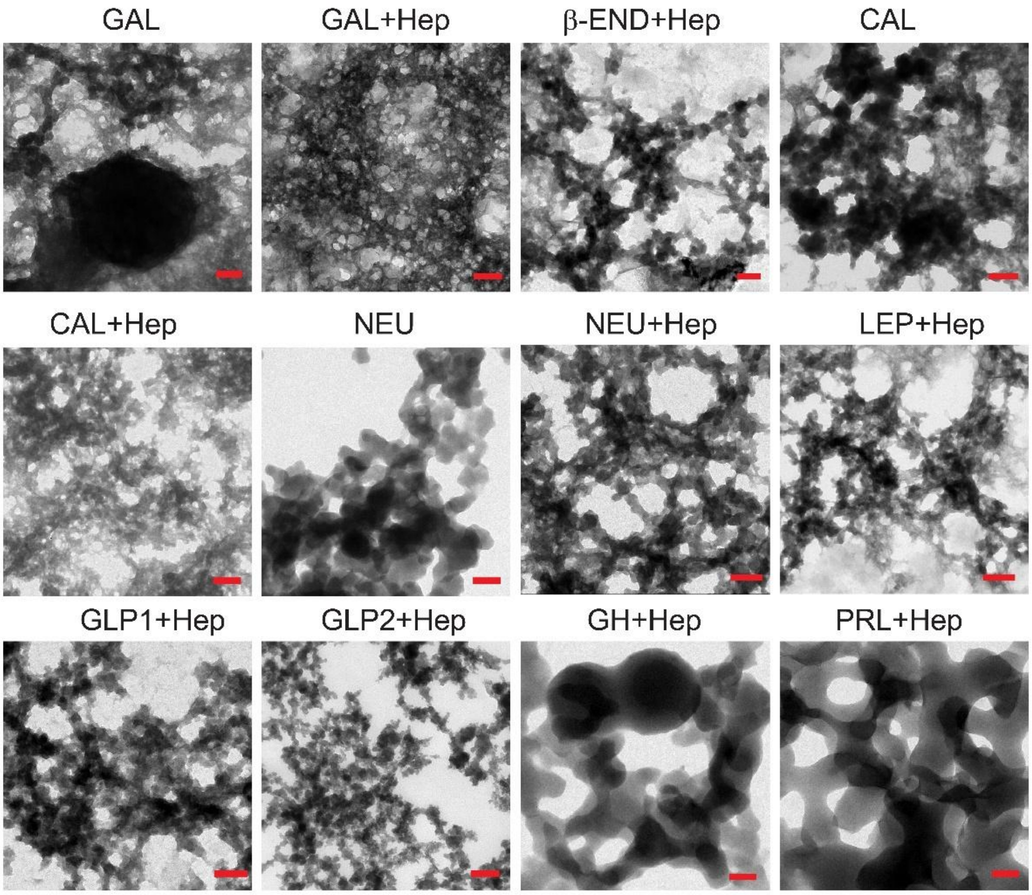


**Figure S7: Morphology of hormone condensates after ageing:** Representative TEM images of different proteins/peptide hormone condensates after 12 h incubation, showing aggregated fibrillar morphology of hormones. Scale bar: 100 nm.

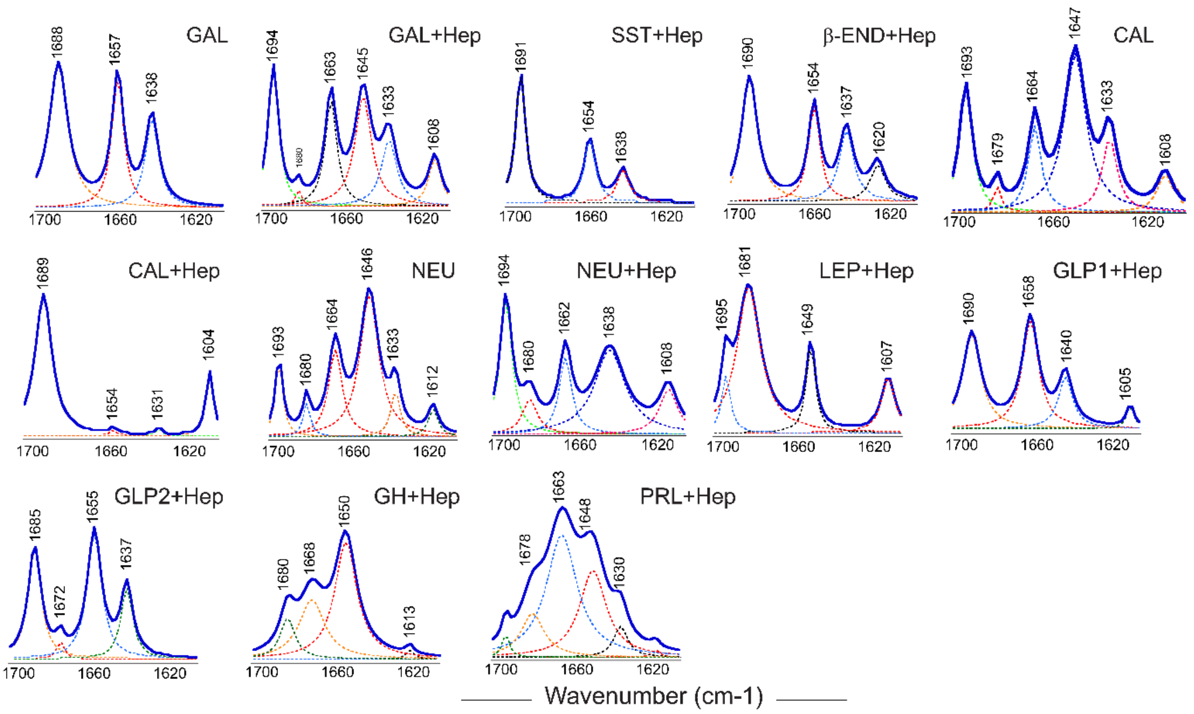


**Figure S8: Secondary structural characterisation of hormone condensates after ageing:** FTIR spectra of different proteins/peptides hormone condensates after 12 h of formation showing amyloid cross-β-sheet conformation specific peaks ~1690 ± 5 $cm^{-1}$ and/or ~1635 ± 5 $cm^{-1}$ (29).

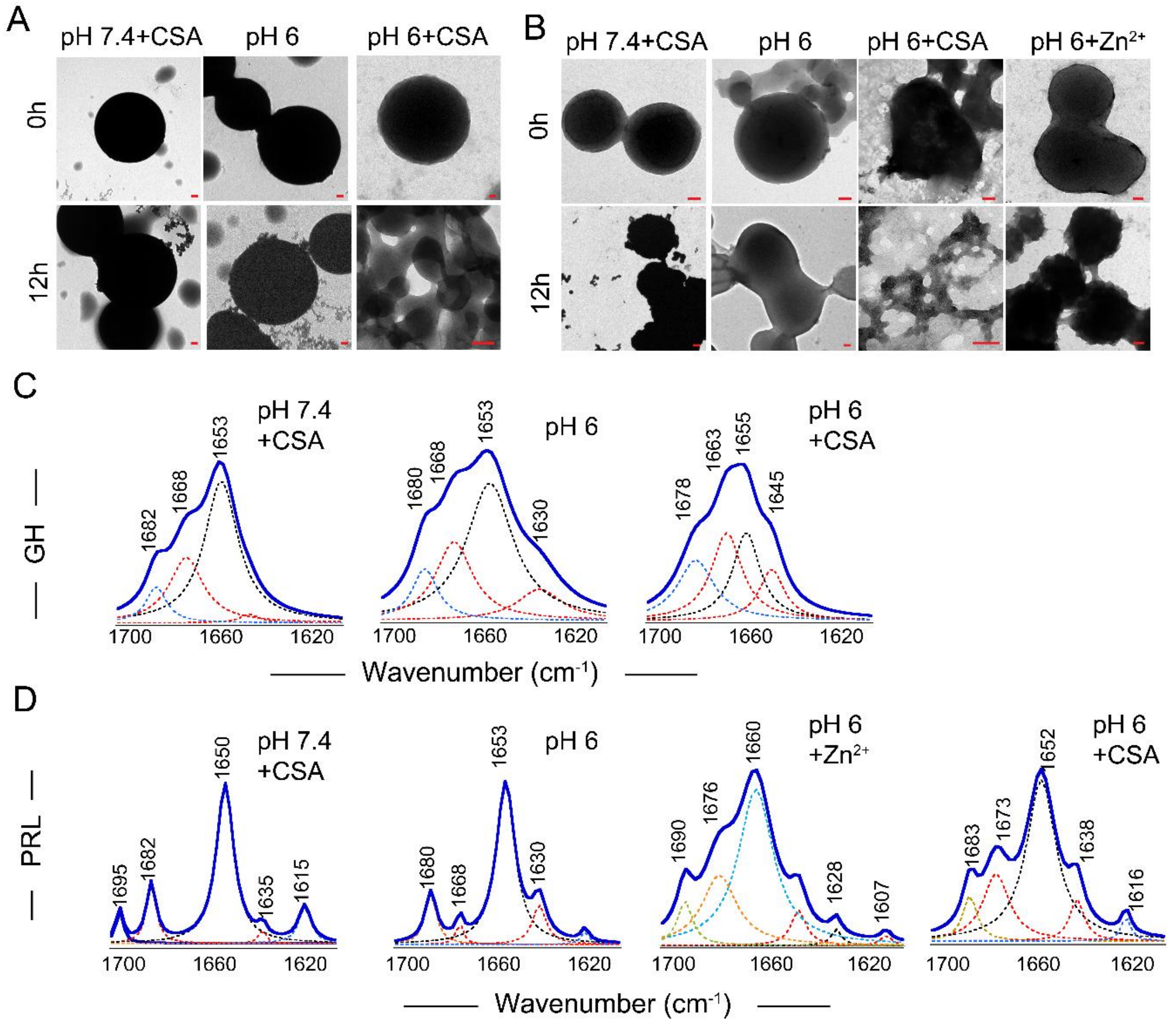


**Figure S9: GH and PRL condensates at various phase separation conditions: (A, B)** Representative TEM images of GH (A) and PRL (B) at various phase separating conditions showing non-fibrillar aggregated morphology after 12h. Scale bar:100 nm. **(C, D)** FTIR spectra of GH (C) and PRL (D) at these phase separating conditions, showing less cross-β-sheet conformation specific peak ~1690 ± 5 $cm^{-1}$ and/or ~1635 ± 5 $cm^{-1}$, at 12 h timepoints after formation (29).

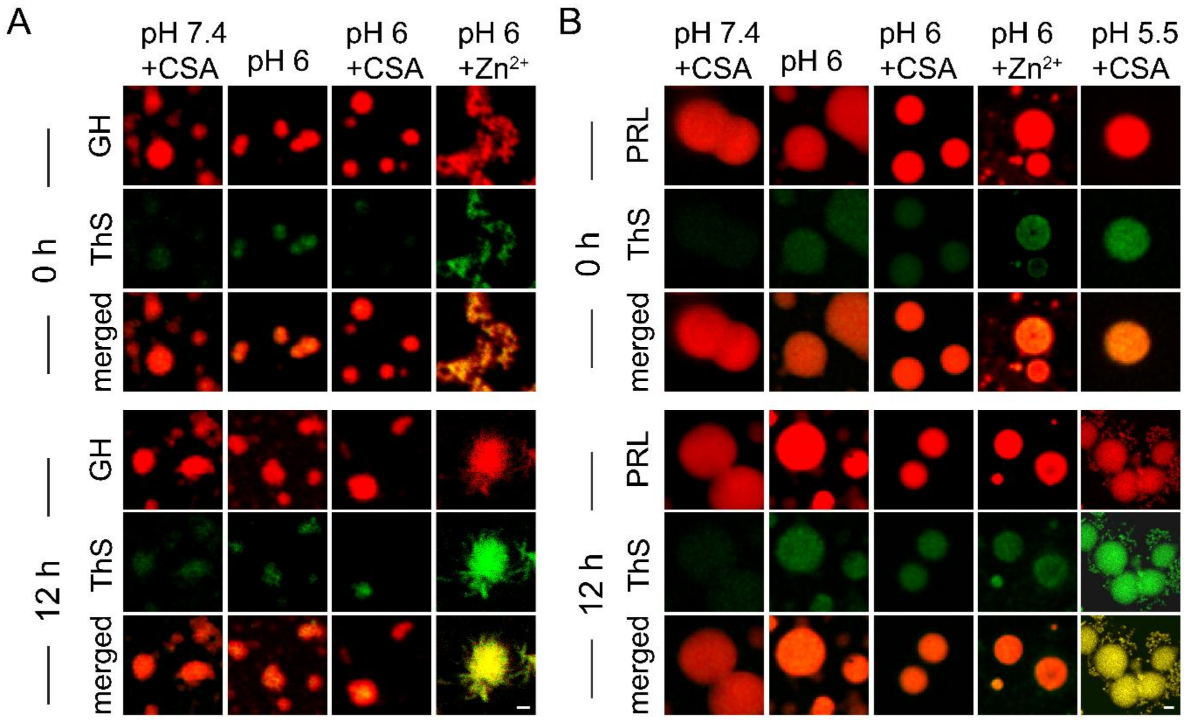


**Figure S10: Time-dependent ThS fluorescence binding of GH/PRL condensates: (A, B) Representative** ThS binding of GH (A) and PRL condensates (B) at different phase-separating conditions immediately after phase separation (0 h) and after 12h of incubation, showing GH in the presence of $Zn^{2+}$ (pH 6) and PRL in the presence of CSA (pH 5.5), showing maximum fluorescence enhancement, indicating their amyloid nature. Star-shaped condensates binding to ThS were observed for GH (pH 6+$Zn^{2+}$), indicating a fibrillar outburst from the condensates. Scale bar: 1μM.

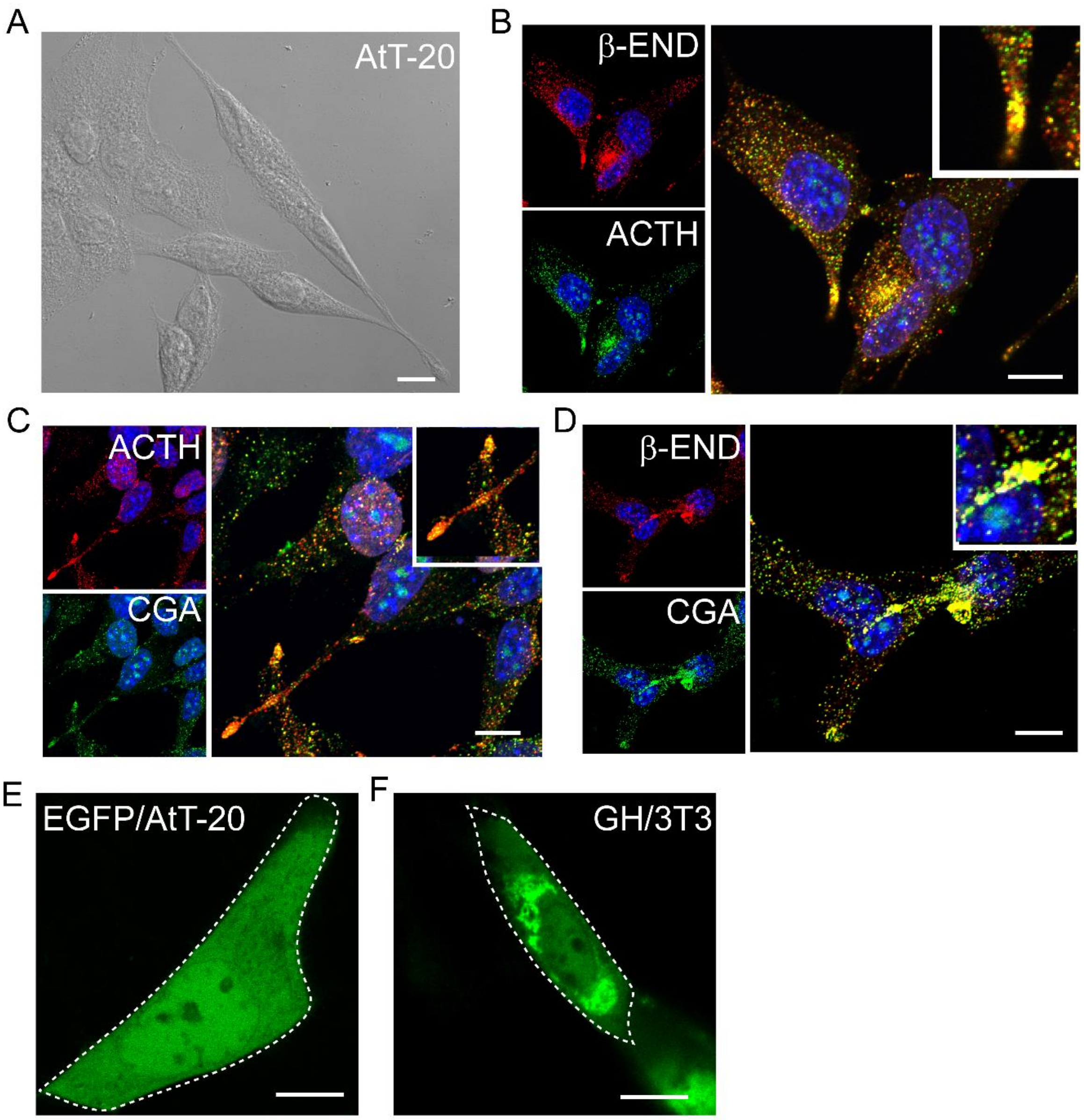


**Figure S11: GH/PRL phase separation in neuroendocrine cell line, AtT-20: (A)** Mouse neuroendocrine cell line AtT-20 used for the study, exhibiting distinct elongated morphology under a phase contrast microscope. **(B-D)** Colocalization of endogenous ACTH and β-END (B); endogenous ACTH and Chromogranin-A (C); and endogenous β-END and Chromogranin-A (D) in AtT20 cells showing the functional characterisation of the cells. (**E-F)** Only EGFP protein expression in AtT-20 cells (E) and GH-EGFP expression in the fibroblast 3T3 cells (F), showing the pan-cellular distribution of the protein with no condensate formation**.** (A-F) Scale bar representing 10 µm.

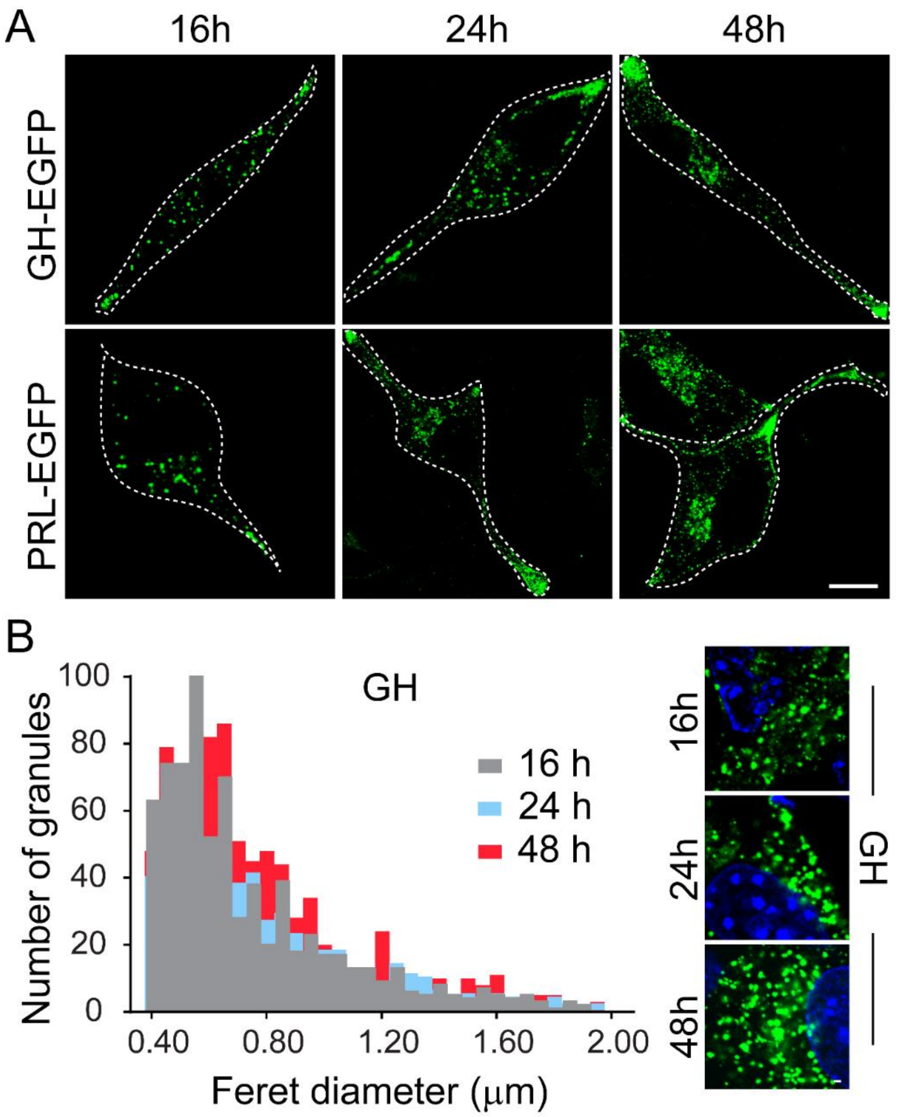


**Figure S12: GH/PRL-EGFP transfected AtT-20 cells showing phase separation over time: (A)** Time-dependent confocal imaging of GH-EGFP (top row) and PRL-EGFP (bottom row) in transfected AtT-20 cells at 16 h, 24 h, and 48 h post-transfection, showing two distinct condensate populations (juxtanuclear and at the cellular tip) post 24 h. Scale bar: 10 μm. **(B)** The histogram analysing the Feret diameter of GH-EGFP condensates at three different timepoints: 16h, 24h, and 48h post-transfection (left). The confocal image of GH condensates at three different time points: 16 h, 24 h, and 48 h. (right). Scale bar:1 μm.

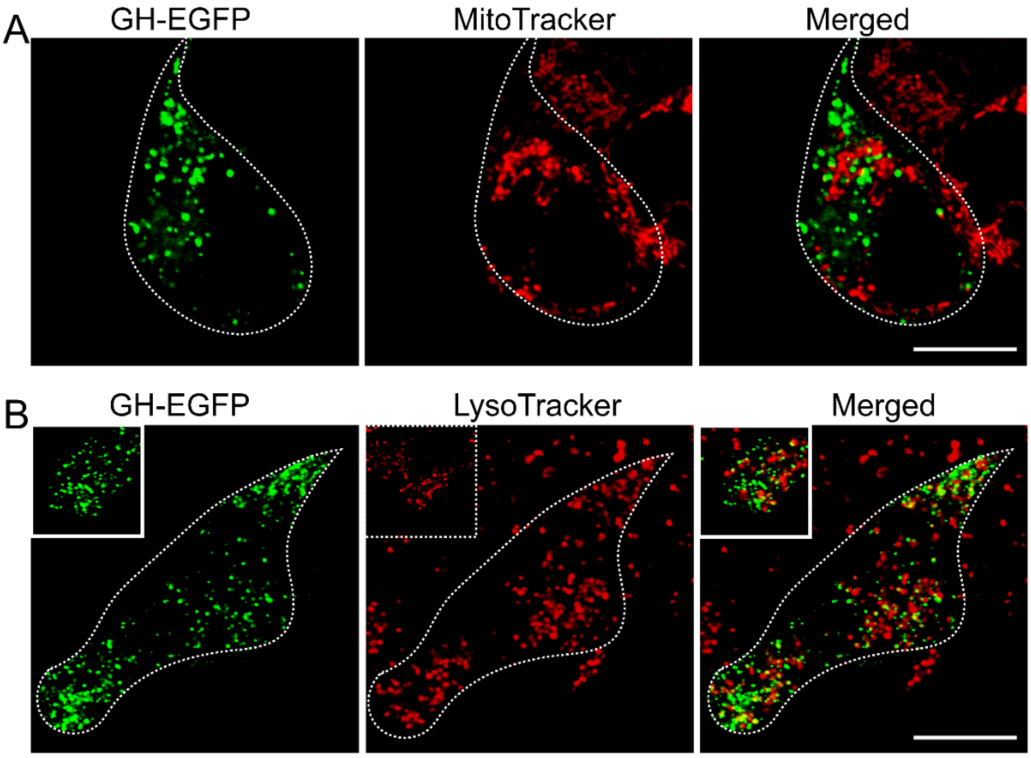


**Figure S13: LysoTracker and MitoTracker staining of Transfected AtT-20 cells: (A-B)** Live cell confocal imaging of GH-transfected AtT-20 cells stained with MitoTracker (red) (A) and LysoTracker (red) (B) dye showing no colocalization with GH granules (green). Scale bar: 10 μm.

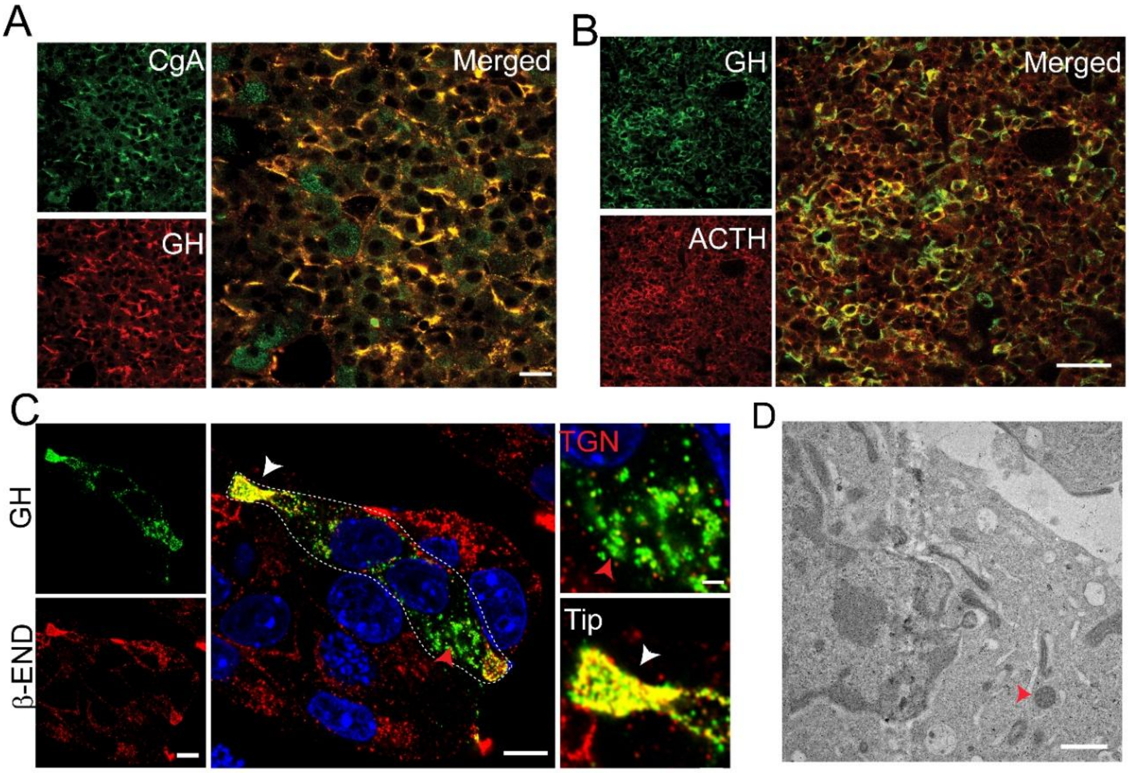


**Figure S14: GH SGs in pituitary tissue and in AtT-20 cells. (A, B)** Rat (Male) pituitary tissue stained with GH and CgA antibody (A), GH and ACTH antibody (B), showing the distribution of these hormones in the anterior pituitary. Scale bar: 50 µm. **(C)** Immunocytochemistry on GH-EGFP-transfected AtT-20 cells with β-END antibody showing granule packaging of GH and endogenous hormones inside cells (left). Scale bar: 10 µm. Zoomed image of TGN and Tip secretory granule population showing the colocalization status of GH hormones with endogenous hormones β-END (right). Scale bar: 2 µm. **(D)** GH-EGFP-transfected AtT-20 cell cross-sectioned at 50 nm through ultramicrotome technology, showing the cell interior with electron-opaque secretory granules (red arrow). Scale bar representing 1µm.

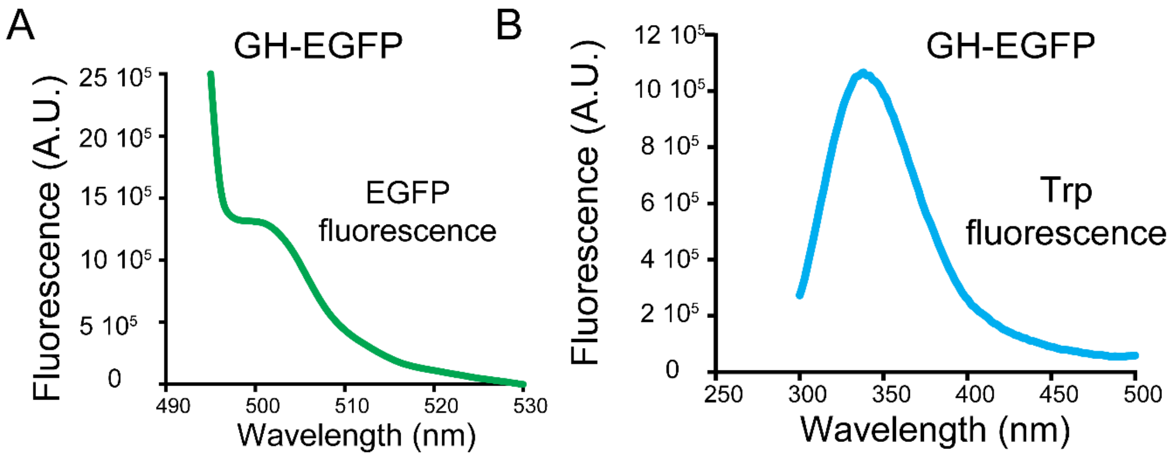


**Figure S15: GH-EGFP SG isolation from AtT-20 cells: (A, B)** GH-EGFP secretory granule-rich cell fraction showing EGFP fluorescence emission spectra (A) at 508 nm when excited at 488 nm and Tryptophan fluorescence emission spectra (B) at 348 nm when excited at 280 nm.

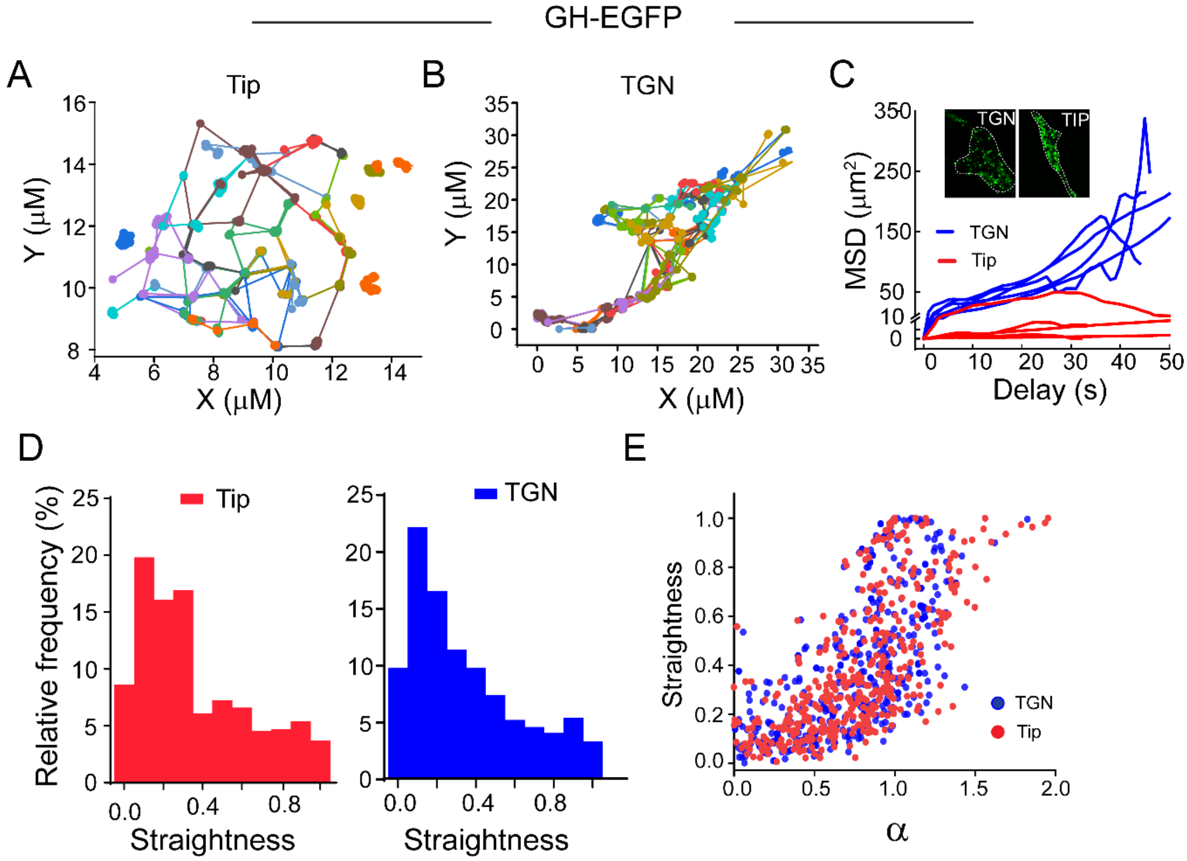


**Figure S16: Single-particle tracking of GH-EGFP condensates/SGs populated at TGN and Tip: (A, B)** The individual GH-EGFP granule trajectory, populated at the Tip (A) and TGN (B) of a representative transfected AtT-20 cell. **(C)** Line plot representing average MSD value of all TGN granules (blue) and Tip granules (red) from a transfection-positive cell, for n=5 cells, showing high dynamicity of condensates at the TGN population. Inset showing a representative magnified image of condensates from the TGN and Tip population of a cell. **(D)** The distribution plot for relative abundance versus the track straightness of 500 condensates/granules populated at TGN and Tip, showing a similar distribution in both locations. **(E)** The plot for granule straightness vs diffusion exponent (α) for 500 granules from TGN and Tip of GH-EGFP transfected AtT-20 cells showing a positive correlation.

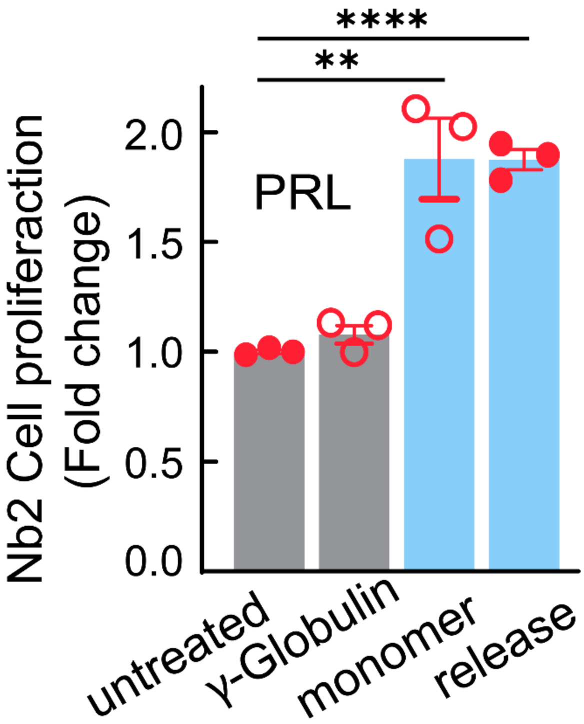


**Figure S17: Functional hormone aggregates in SGs:** Fold change in Nb2 cell proliferation with the released PRL hormone from amyloid-rich condensates, along with unrelated protein γ-Globulin. The monomeric PRL was used as a positive control, and untreated cells were used as a negative control. The statistical significance was calculated using a two-tailed unpaired t-test [***P*<0.009, *****P* < 0.0001 with a 95% confidence interval].

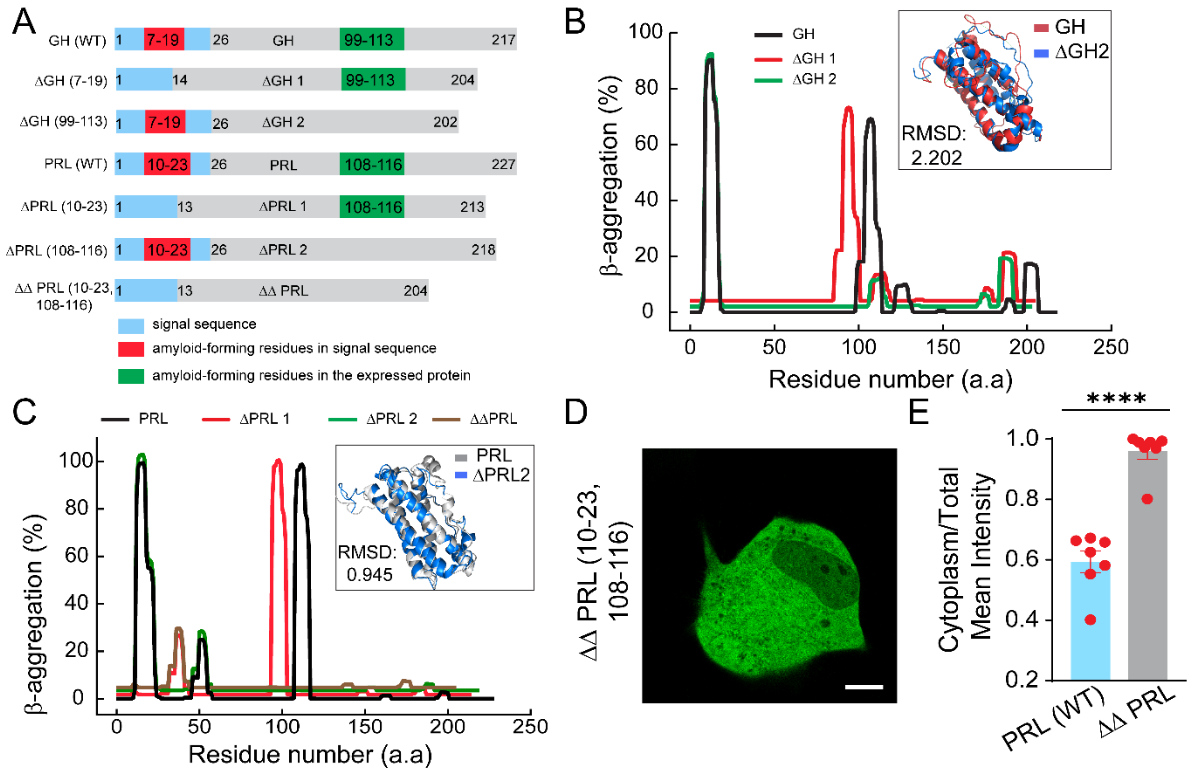


**Figure S18: Role of amyloidogenic sequence in GH/PRL condensate formation, liquid-to-solid transition and in SG biogenesis: (A)** Schematic representation depicting all the deletion mutant constructs of GH and PRL with sequential deletion of the amyloidogenic regions. **(B, C)** TANGO analysis of full-length GH (B) and PRL (C), along with their deletion mutants, showing the removal of the amyloid-scoring amino acid regions in the deletion mutants. The inset showing the superimposed AlphaFold structure of ΔGH2 and ΔPRL2 with the respective WT protein structure, with the RMSD scores. **(D, E)** Confocal live cell image (D) and corresponding quantification of Cytoplasm/Total Mean Intensity (n=7 cells) (E) showing pan cellular protein distribution without any SG formation upon transfection with ΔΔPRL (10-23, 108-116) variant. Scale bar 10 µm.

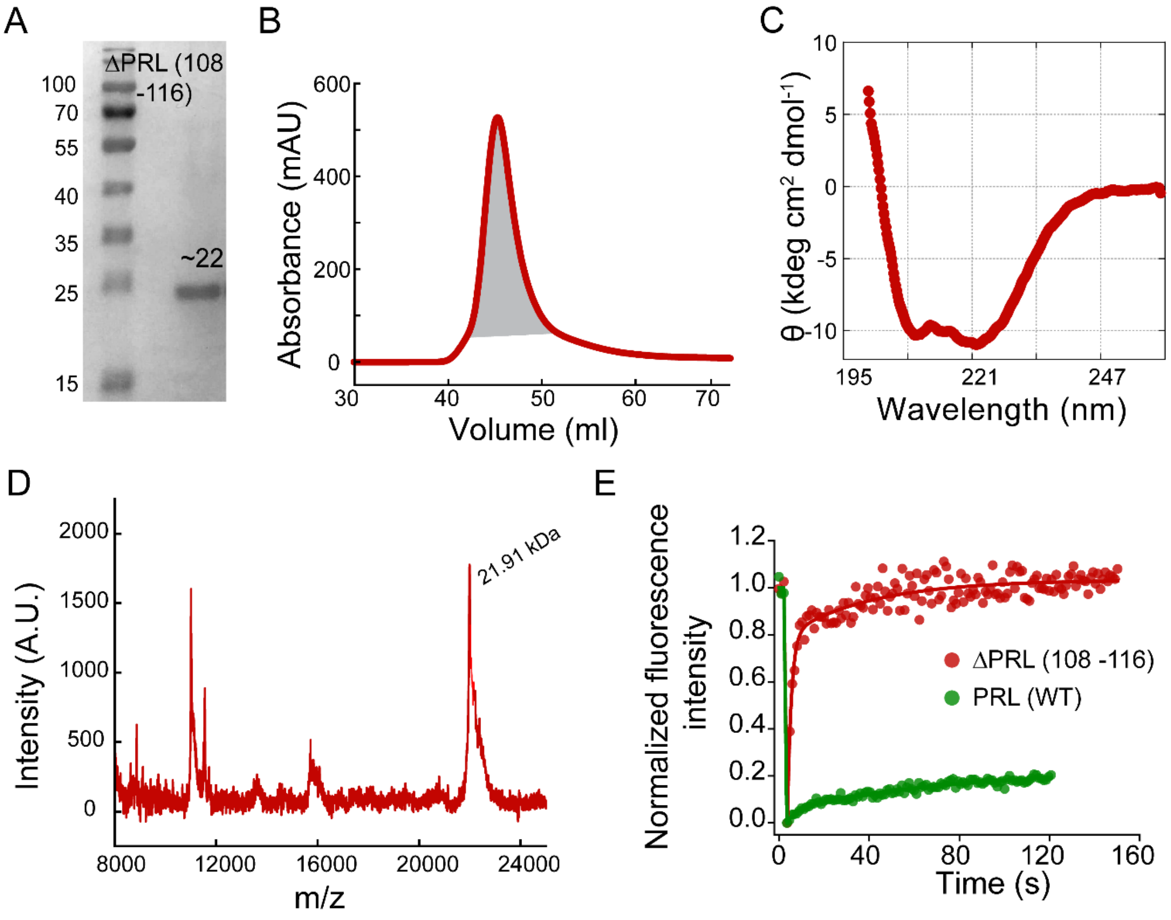


**Figure S19: Characterisation and liquid-like phase separation of ΔPRL (108-116): (A-D)** The purified ΔPRL (108-116) proteins were characterised via SDS Page, showing a single band at a molecular weight of ~22 kDa (A) and single peak in size exclusion chromatogram (B), α-helical structure by CD spectroscopy (C) and molecular mass by MALDI TOF mass-spectrometry (D), confirming the purified protein. **(E)** Normalized fluorescence intensity of ΔPRL (108-116) and WT PRL condensate (immediately formed) at pH 5.5 in the presence of CSA, showing liquid-like high FRAP recovery for the mutant protein, while WT PRL showed very low FRAP recovery.

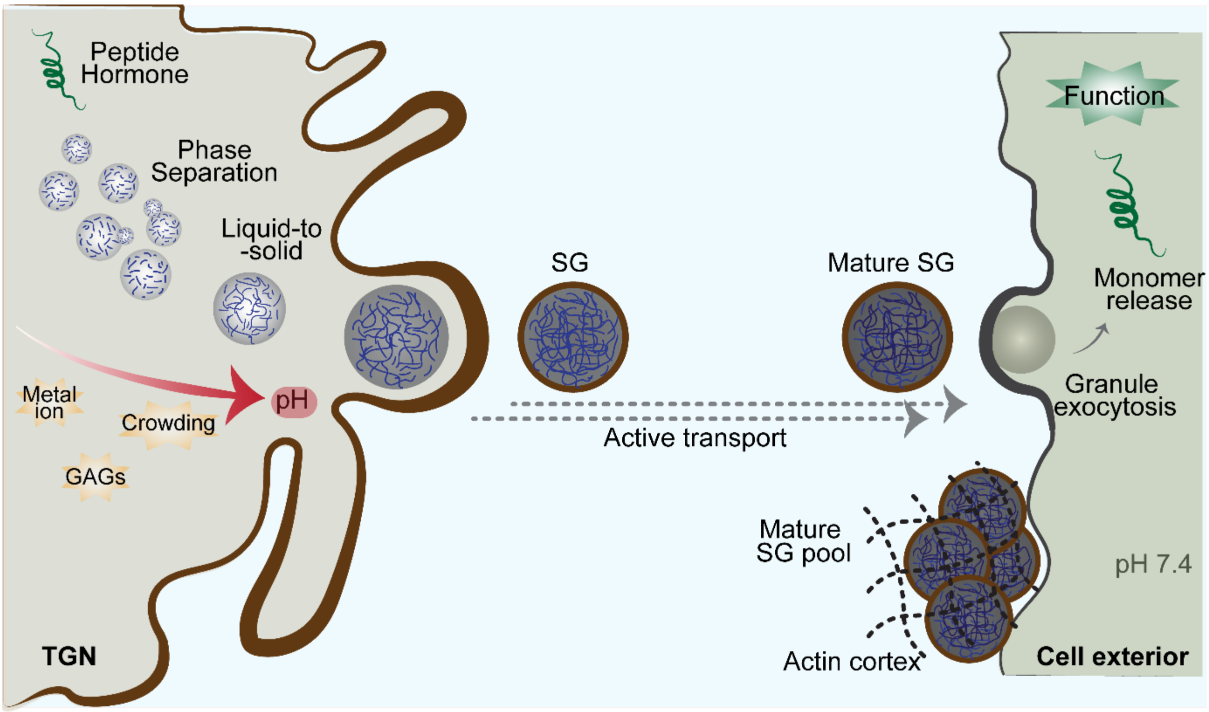


**Figure S20: Phase separation of hormones at TGN driving functional aggregation as the rapid storage mechanism of hormones during granule biogenesis:** Schematic representation of the proposed molecular mechanism of secretory granule biogenesis through hormone phase separation. The TGN microenvironment (low pH, metal ions, GAGs, and molecular crowding) promotes both phase separation and a rapid liquid-to-solid transition of protein and peptide hormones, thereby concentrating and segregating them from the surrounding milieu and forming a nascent condensate/SG pool at the TGN. Following membrane-budding, these SGs are actively transported through the microtubule network to the distal cell periphery to be stored as a mature secretory granule pool, tethered by the actin cortex. During the transport, SG maturation also takes place, where the membrane-wrapped solid-like condensates turn into functional amyloid over time, forming the dense core of the mature secretory granules. Upon regulated exocytosis, these amyloid aggregates revert to native, soluble hormone in the extracellular environment (pH 7.4 and dilution), resulting in the burst release of bioactive proteins.

**Supplementary Table**

**Table S1. Details of protein/peptide hormones:** Complete list of all the hormones under study, including their amino acid sequence, location, secondary structure, amino acid length, and molecular weight.

| No | Name | Abbreviation | PDB ID | Peptide Sequence | A.A | M.W. | Abundance/ isolated |
|---|---|---|---|---|---|---|---|
| 1 | Galanin | GAL | 8DJ4 | GWTLNSAGYL LGPHAVGNHR SFSDKNGLTS | 30 | 3157 | Colon and Pituitary |
| 2 | Somatostatin-14 | SST | 2MI1 | AGCKNFFWKT FTSC (disulfide bond) | 14 | 1639 | Pancreas |
| 3 | β-Endorphin | β-END | 8F7Q_2 | YGGFMTSEKS QTPLVTLFKN AIIKNAYKKG E | 31 | 3465 | Pituitary |
| 4 | Salmon Calcitonin | CAL | 2GLH | CSNLSTCVLG KLSQELHKLQ TYPRTNTGSG TP | 32 | 3435 | Thyroid |
| 5 | Glucagon-like Peptide 2 | GLP2 | 2L63 | HADGSFSDEM NTILDNLAAR DFINWLIQTK ITD | 33 | 3766 | Pancreas |
| 6 | Glucagon-like Peptide 1 | GLP1 | - | HDEFERHAEG TFTSDVSSYL EGQAAKEFIA WLVKGRG | 37 | 4170 | Pancreas |
| 7 | Neurotensin | NEU | 2LNE_A | LYENKPRRPY IL | 12 | 1562 | GI tract and CNS |
| 8 | Leptin | LEP | 8X80_D | MHWGTLCGFLWLWPYLFYVQAVPI QKVQDDTKTLIKTIVTRINDISHTQS VSSKQKVTGLDFIPGLHPILTLSKMD QTLAVYQQILTSMPSRNVIQISNDLE NLRDLLHVLAFSKSCHLPWASGLET LDSLGGVLEASGYSTEVVALSRLQGS LQDMLWQ LDLSPGC | 167 | 18640 | White adipose tissue |
| 9 | Growth Hormone | GH | 1HGU | FPTIPLSRLFDNAMLRAHRLHQLAFD TYQEFEEAYIPKEQKYSFLQNPQTSL CFSESIPTPSNREETQQKSNLELLRISL LLIQSWLEPVQFLRSVFANSLVYGAS DSNVYDLLKDLEEGIQTLMGRLEDG SPRTGQIFKQTYSKFDTNSHNDDALL | 191 | 22129 | Pituitary |

| | | | | | | | |
|---|---|---|---|---|---|---|---|
| | | | | KNYGLLYCFRKDMDKVETFLRIVQC RSVEGSCGF | | | |
| 10 | Prolactin | PRL | 1RW5 | LPICPGGAARCQVTLRDLFDRAVVLS HYIHNLSSEMFSEFDKRYTHGRGFIT KAINSCHTSSLATPEDKEQAQQMNQ KDFLSLIVSILRSWNEPLYHLVTEVR GMQEAPEAILSKAVEIEEQTKRLLEG MELIVSQVHPETKENEIYPVWSGLPS LQMADEESRLSAYYNLLHCLRRDSH KIDNYLKLLKCRIIHNNNC | 199 | 22898 | Pituitary |

**Table S2. Summary of protein/peptide phase-separation conditions.** Summary of all phase-separation conditions examined for protein and peptide hormones, along with the material properties of the resulting condensates immediately after formation. (*Note: 500 µM heparin caused immediate dissolution of SST condensates.)

| Protein/ Peptides | Concentration | Phase-separation Conditions | Condensates formation | Fusion/Wetting (0h) |
|---|---|---|---|---|
| GAL | 2 mg/mL | pH 6 | Yes | Yes |
| GAL | 2 mg/mL | pH 6+ 500 µM Hep | Yes | No |
| SST | 2 mg/mL | pH 6+ 50 µM Hep* | Yes | Yes |
| β-END | 2 mg/mL | pH 6+ 500 µM Hep+ 5% PEG-8000 | Yes | No |
| CAL | 2 mg/mL | pH 6 | Yes | Yes |
| CAL | 2 mg/mL | pH 6+ 500 µM Hep | Yes | No |
| GLP2 | 2 mg/mL | pH 6+ 500 µM Hep+ 5% PEG-8000 | Yes | No |
| GLP1 | 2 mg/mL | pH 6+ 500 µM Hep+ 5% PEG-8000 | Yes | No |
| LEP | 2 mg/mL | pH 6+ 500 µM Hep | Yes | No |
| NEU | 2 mg/mL | pH 6 | Yes | Yes |
| NEU | 2 mg/mL | pH 6+ 500 µM Hep | Yes | No |
| GH | 2 mg/mL | pH 6+ 500 µM Hep | Yes | No |
| GH | 100 µM- 1.5 mM | pH 7.4+ CSA/CSB/Hep (100 µM- 1.0 mM) | Yes | Yes |
| GH | 250 µM | pH 6 | Yes | No |
| GH | 100 µM | pH 6+ 100 µM CSA | Yes | No |
| GH | 100 µM | pH 6 +100 µM $ZnSO_4$ | Yes | No |
| PRL | 2 mg/mL | pH 6+ 500 µM Hep | Yes | No |
| PRL | 50 µM- 1.2 mM | pH 7.4+ CSA/CSB/Hep (50 µM- 500 µM) | Yes | Yes |
| PRL | 200 µM | pH 6 | Yes | No |
| PRL | 100 µM | pH 6+ 50 µM CSA | Yes | No |
| PRL | 100 µM | pH 5.5+ 50 µM CSA | Yes | No |
| PRL | 100 µM | pH 6 +50 µM $ZnSO_4$ | Yes | No |

**Table S3. List of all GH/PRL deletion variants:** The list of all GH and PRL variants with amino acid details for sequential deletion of the amyloid sequence from the GH and PRL amino acid sequence.

| GH/PRL variant | Abbreviation | Deleted sequence |
|---|---|---|
| GH WT | GH | - |
| ΔGH (7-19) | ΔGH1 | TSLLLAFGLLCLP |
| ΔGH (99-113) | ΔGH2 | LELLRISLLLIQSWL |
| PRL WT | PRL | - |
| ΔPRL (10-23) | ΔPRL1 | GSLLLLLVSNLLLC |
| ΔPRL (108-116) | ΔPRL2 | FLSLIVSIL |
| ΔΔPRL (10-23, 108-116) | ΔΔPRL | GSLLLLLVSNLLLC and FLSLIVSIL |

**Supplementary Video Legends**

**Supplementary Video 1:** Liquid-like fusion property of GH (pH 7.4+CSA), in-vitro condensates. Scale bar: 1 µm.

**Supplementary Video 2:** Liquid-like fusion property of NEU (pH 6) in-vitro condensates. Scale bar: 1 µm.

**Supplementary Video 3:** Time-lapse video showing fusion event at the TGN population of AtT-20 cells. Scale bar: 1 µm.

**Supplementary Video 4:** Lattice light-sheet microscopy imaging and corresponding time-lapse video of highly dynamic GH-EGFP condensates accumulated at the TGN of the AtT-20 cells. 5 µm grid.

**Supplementary Video 5:** Lattice light-sheet microscopy imaging and corresponding time-lapse video of the GH-EGFP condensates accumulated at the Tip of the AtT-20 cells. Scale bar: 5 µm grid.
